# Genomic signatures of terrestrial adaptation in air-breathing catfishes (Clariidae)

**DOI:** 10.1101/2025.03.20.644309

**Authors:** Gopi Krishnan, Shivakumara Manu, Sreenivasu Ara, Rajeev Raghavan, Govindhaswamy Umapathy

## Abstract

Air-breathing catfishes of the family Clariidae exhibit extraordinary adaptations that enable them to survive outside water for extended periods, yet the genetic and genomic basis of these adaptations remain poorly understood. To study these adaptations, we sequenced and assembled two high-quality genomes of two clariid species, *Clarias gariepinus* and *Clarias dussumieri* and compare them with previously available genomes of 23 catfish species across nine families. By reconstructing the whole-genome phylogeny and examining patterns of positive selection and gene family evolution, we found unique signatures associated with terrestrial adaptation in clariids. Our analysis revealed that a high proportion of genes were positively selected in clariids, that play critical roles in hypoxia tolerance, thermoregulation, metabolism, and DNA repair, which are key traits for terrestrial adaptation. Additionally, we observed significant expansions in gene families, including Myoglobin (involved in oxygen transport), immunity-related genes, and xenobiotic degradation pathways, highlighting their importance in environmental resilience and detoxification. Together, these findings provide a comprehensive understanding of the genomic changes facilitating the terrestrial adaptation of clariids. This study also highlights the contribution of genome evolution to their resilience, adaptability to novel environments, and invasiveness, offering valuable insights into the genetic basis of ecological niche diversification.

## Introduction

Ecological transitions of species to novel habitats have long been a subject of interest in evolution and ecology, as they offer insights into the mechanisms driving adaptation. Such transitions have occurred across diverse lineages throughout evolutionary history. For instance, some of the key events include the origin of tetrapods (e.g. Tiktaalik) around 385–359 million years ago (Daeschler et al. 2006) and a relatively recent transition of cetaceans ∼50 MYA (Fordyce 2018). Investigating these ecological transitions allows us to explore the evolutionary mechanisms facilitating such adaptations and the genetic and genomic bases underlying them. Yuan et al. (2021) suggest that while ancient transitions may obscure the genomic basis of these adaptations due to extended evolutionary periods, more recent transitions may offer precise insights.

One such fascinating example of ecological transition is the semi-terrestrialization of fishes, which has occurred multiple times across different families throughout the evolutionary past. Some of the significant events include the evolution of lungfishes (Liem 1988), mudskippers (Jaafar & Murdy 2017), climbing perches (Davenport and Matin 1990) and notably that of, air-breathing catfishes of the family Clariidae (Bruton 1979). While lungfishes have developed true lungs, mudskippers and climbing perches have acquired phenotypic modifications such as modified pectoral fins, ammonia tolerance, aerial respiration etc. (Davenport et al. 1990, You et al. 2014) allowing them to survive on land for short periods.

Members of the family Clariidae (henceforth Clariids) possess a unique adaptation in the form of accessory respiratory organs, which are modified gill structures that enhance oxygen exchange (Chandra and Banerjee 2003). As obligate breathers, these organs account for 85% of their oxygen uptake (Maina and Maloiy 1986), rendering their gills ineffective in well-aerated and deep waters. This adaptation has allowed clariids to colonize land and become popular in aquaculture due to their low maintenance requirements. In fact, they are one of the major aquaculture species worldwide (Na-Nakorn and Brummet 2009). Their ability to survive in poorly-oxygenated & polluted waters, heat tolerance, and extended burrowing periods have also contributed to their invasiveness (Cambray 2003; Mahapatra and Mohanty 2023; Singh et al. 2015). As members of the second most diverse group of fishes, the catfishes (Siluriformes) (Diogo 2003), clariids provide an excellent model for investigating the genomic evolution and genetic basis of these unique traits and terrestrial adaptation.

While extensive studies have explored the morphological and anatomical features of clariid fishes, the genomic and molecular basis for their terrestrial adaptation remains underexplored (Li et al. 2018b; Kushwaha et al. 2021; Nguinkal et al. 2024). Specifically, the genomic mechanisms underlying adaptation to novel environments have received little attention. To address this knowledge gap, we carried out a comparative genomics study including clariids and other catfish species. We sequenced and annotated the genomes of two clariid species (the widely distributed invasive African Catfish, *Clarias gariepinus* and the peninsular Indian endemic catfish, *Clarias dussumieri*) and included the published genomes of 23 other catfish species, covering nine families, using a robust bioinformatics pipeline. We hypothesized that clariids would exhibit specific genomic signatures of strong selection and gene family expansions related to traits supporting terrestrial adaptation. To investigate this, we first constructed a whole-genome phylogeny and estimated the time these lineages diverged. Leveraging this phylogenetic framework and streamlined data, we then undertook a genome-wide screening for positively selected genes in clariids. Finally, we examined gene family expansions and the enrichment of gene ontology terms associated with terrestrial adaptation.

## Methods

### 1. Genome sequencing and assembly

We generated two high-quality draft genome assemblies for *Clarias gariepinus* and *Clarias dussumieri* using a hybrid approach that combined long-read data from Oxford Nanopore Technology (ONT) and short-read data from the Illumina NovaSeq platform. The genome was initially assembled de novo using ONT long reads with Canu v2.0 (Koren et al. 2017). The Illumina short reads were pre-processed using fastp v0.23.2 (Chen et al. 2018), which we then used to polish the draft assembly using the POLCA pipeline in Masurca v4.0.8 (Zimin et al. 2013). To assess genome completeness and quality, we employed BUSCO (Benchmarking Universal Single-Copy Orthologs) (Manni et al. 2021) with the Actinopterygii dataset (OrthoDB v10), which comprises 3,640 conserved genes. For the comparative genomics workflow, we obtained the genomes of 23 catfish species (Order: Siluriformes) belonging to nine families from NCBI GenBank (see supplementary file, Table 1 for the complete list and statistics). These genomes had an N50 value of at least 10Kb and Busco completeness of at least 85%. Additionally, we included the genomes of *Danio rerio* (zebrafish) and *Oreochromis niloticus* (Nile tilapia) as outgroups.

### 2. Gene prediction and functional annotation

To minimize potential technical artifacts arising from the use of different tools across species, we applied the same bioinformatics pipeline for repeat masking, gene prediction and functional annotation to all 27 genomes, thus providing a solid foundational dataset. For repeat content characterization, we first built a custom repeat library for each species using RepeatModeler v2.0.5 (Flynn et al. 2020), supplied with curated Dfam database. The respective custom libraries were then used to predict repeat content and soft-mask the genomes with RepeatMasker v4.1.5 (Smit et al. 2013). The identified repeat elements were classified into categories including Retroelements (SINEs, Penelope, LINEs, and LTR elements), DNA transposons, Rolling-circle elements, Small RNAs, Satellites, and Simple repeats.

We performed gene prediction for the soft-masked genomes using the GALBA v1.0.8 pipeline (Brůna et al. 2023), a fully automated tool specifically designed for gene prediction in large genomes lacking RNA-Seq data. We supplemented the pipeline with 7.5 million Actinopterygii protein sequences downloaded from the UniProt database. After gene prediction, sequences unsupported by evidence were removed, and the predicted set was pruned by retaining only the longest isoforms. The final gene prediction sets were evaluated for completeness using BUSCO against the Actinopterygii dataset (OrthoDB v10). Functional annotation of the predicted protein sequences was performed using eggNOG-mapper v2.1.12 (Cantalapiedra et al. 2021). Additionally, Gene Ontology (GO) terms were assigned to the proteins using the FANTASIA pipeline (Martínez-Redondo et al. 2024).

### 3. Phylogenetic tree construction

We assigned orthology for the predicted protein sequence sets of the 27 species using Orthofinder v2.5.5 (Emms and Kelly 2019), retrieving both single-copy (SCO) and multi-copy orthologous sequences. The SCOs were aligned at the amino acid level using MAFFT. Corresponding nucleotide sequences for the aligned SCOs were extracted from the coding sequence files generated by GALBA and aligned using the codon-aware aligner MACSE (Ranwez et al. 2011). The resulting nucleotide alignments were trimmed with TrimAl v1.4.1 (Capella-Gutiérrez et al. 2009) to remove gaps. The trimmed alignments were concatenated using AMAS (Borowiec et al. 2016) and was then used as input for maximum likelihood based phylogenetic inference in IQ-TREE v2.3.2 (Minh et al. 2020), under the GTR model of sequence evolution, automatically selected as the best fit by ModelFinder (Kalyaanamoorthy et al. 2017). The resulting species tree was visualized using FigTree v1.4.4.

### 4. Divergence time estimation

We estimated species divergence times using the MCMCTree module of the PAML package v4.10.7 (Yang et al. 2007). First, we extracted the fourfold degenerate third codon positions (4DTV regions) from the concatenated gene alignment. The species tree generated by IQ-TREE was provided as the guide tree along with calibration points for seven nodes, for divergence time estimation. For every single chain, we ran 20 million iterations with 100,000 burn-in steps and sampled every 20,000 iterations. To ensure robustness, we performed six independent chains for both prior and posterior analyses. We also assessed the convergence among the chains for both prior and posterior distributions. The resulting ultrametric tree was visualized using FigTree v1.4.4 and subsequently utilized for gene family evolution analyses.

### 5. Positive selection analysis

To identify genes and specific sites under positive selection in clariids, we used the CODEML module of the PAML package v4.10.7 on the nucleotide alignments of single-copy orthologs (SCOs). The Clariidae lineage was designated as the foreground, while all other lineages served as the background. CODEML was run under the branch-site model for each alignment separately, followed by a second run using the null model with a fixed omega value (fix_omega = 1, omega = 1). A likelihood ratio test (LRT) was performed by comparing the log-likelihood values between the null and alternative models to assess the statistical significance of positive selection. Positively selected genes (PSGs) were filtered by having p-value <0.05 from the LRT test, and requiring at least one site with a posterior probability > 0.95, as identified through Bayes Empirical Bayes (BEB) analysis. Gene functions were annotated using results from eggNOG-mapper. For comparison, the same analysis was conducted on four additional lineages: Bagridae, Ictaluridae, Pangasidae, and Siluridae. Lineages consisting of only a single species were not analyzed as foreground branches due to limited comparative power, but were always included in the background set in all analyses.

### 6. Gene family evolution analysis

We analyzed changes in gene family sizes specific to the Clariidae lineage using CAFE5 (Mendes et al. 2020). The gene count file generated from the Orthofinder run was supplied as input, along with the ultrametric phylogenetic tree derived from divergence time estimation. CAFE5 was then used to identify significant expansions or contractions in gene families. First, we filtered out gene families that had evolved significantly (adjusted p-value < 0.05, Benjamini &1Hochberg’s method). We then further refined the dataset by selecting orthogroups that exhibited expansions specifically in Clariidae. To identify Gene Ontology (GO) terms enriched in the expanded gene families of clariids, we conducted GO enrichment analysis using the TopGO R package (Alexa and Rahnenführer 2009).

See the ‘Methods’ section in the Supplementary file for the detailed methods, packages, parameters, and scripts used in various methodological approaches.

## Results and discussion

### 1. Genome assembly and annotation

Through long-read sequencing, we obtained 32x coverage data for *Clarias gariepinus* and 42x coverage data for *Clarias dussumieri*. *De novo* assembly of this data resulted in a 1.18 Gb assembly comprising 3,320 contigs with a N50 of 2.35 Mb for *C. gariepinus*, and a 1.08 Gb assembly comprising 2,416 contigs with a N50 of 3.96 Mb for *C. dussumieri* (Fig. 1a). The base accuracy of the genomes was above 99.5% after polishing them with about 100x coverage of high-quality short reads. BUSCO genome assessment also showed that both genomes had about 99% completeness. measured with the Actinopterygii single-copy gene catalog.

**Fig. 1:**
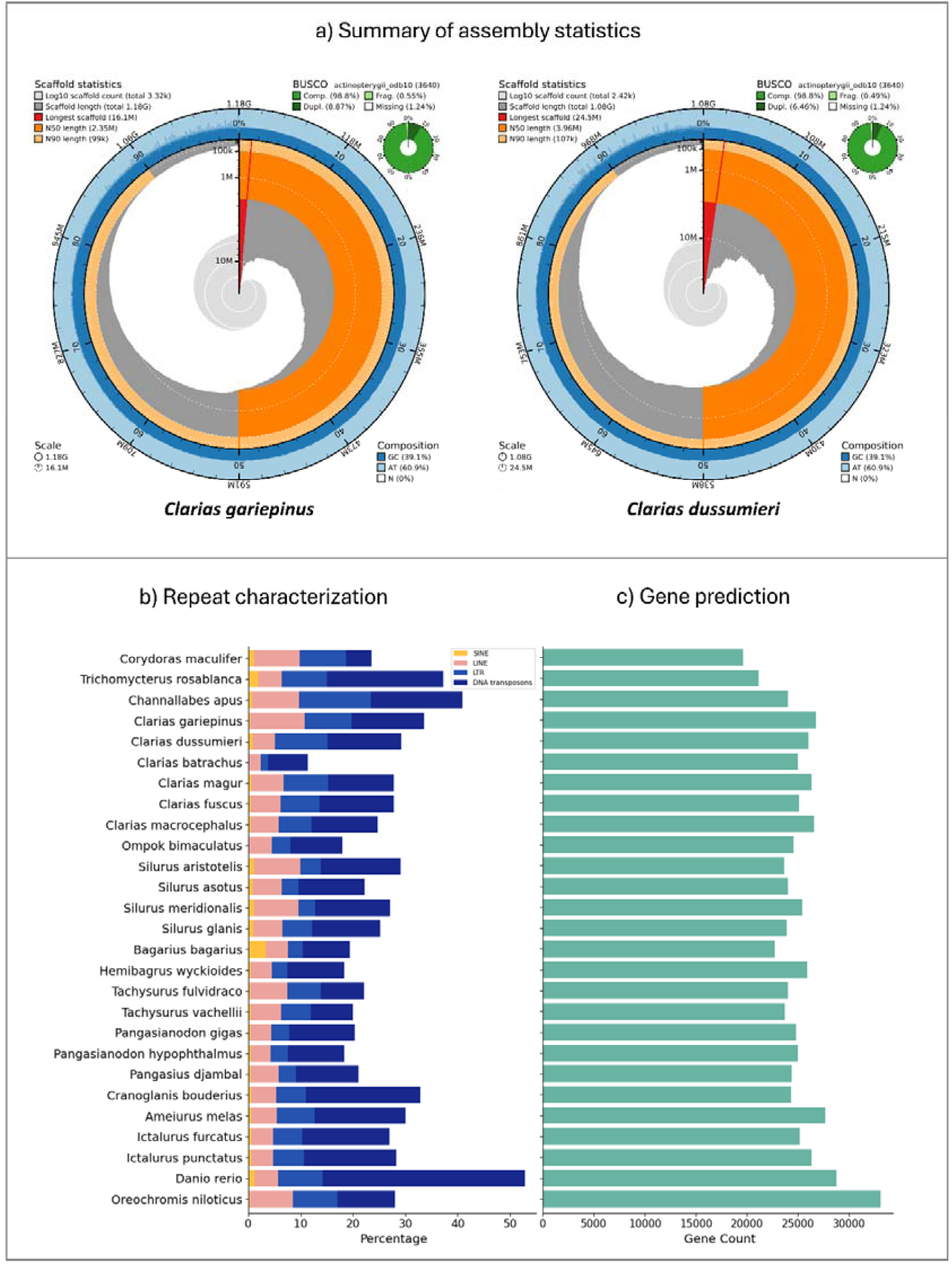
a) Summary of the genome assembly statistics for *Clarias gariepinus*and *Clarias dussumieri*. b) Percentage of genome covered by major categories of Repeat contents across catfish species and outgroups. c) Predicted gene count for catfish and outgroup species.

Repeat sequence annotation of the two assembled genomes, 23 other catfish genomes, and two outgroups revealed a widespread variation in the repeat content across species (Fig. 1b). In total, the genome of *T. rosablanca* was annotated with 55.03% repeat content (highest among the catfishes), whereas the lowest was observed in *B. bagarius* with 29.67% (See supplementary table 2). Gene prediction using the GALBA pipeline and further filtering yielded the expected gene count across species. Among catfishes, the number of predicted genes ranged from 19,629 to 27,701 (Fig. 1c), which closely aligns with previously reported data (Waldbieser et al. 2023; Xu et al. 2022; Dhar et al. 2019; Li et al. 2018b). BUSCO analysis of the predicted proteome revealed that the average completeness is around 94% (±3.6%) (See supplementary Table 3).

In total, 678,196 genes were annotated across 27 genomes. About 98.4% of this set of genes were assigned to orthogroups to enable comparison of orthologous genes across species. We identified a total of 28,223 orthogroups, with the mean orthogroup size being 23.6. There were 10,666 orthogroups that contained sequences from all species. Among these, 5,253 were single-copy orthologs, of which 5,241 were used for the subsequent phylogenetic analysis after alignment and trimming.

### 2. Phylogeny and divergence time estimation

After trimming and concatenating the nucleotide sequences of 5241 single-copy orthologs, we obtained a gene alignment of 6,449,817 base pairs, with 15,723 partitions. The inferred species tree was re-rooted with the outgroup *O. niloticus*, and was well-supported with bootstrap values of 100% for all nodes, except one, which had 98% (Supplementary figure 1). The tree topology is consistent with the geographical distribution of the species, and the species clustered within their families as expected. While the species from the two South American catfish families (Callichthyidae and Trichomycteridae) were recovered together and separated from the rest, the other seven families from Africa & Asia (Clariidae, Siluridae, Sisoridae, Bagridae, Pangasiidae), and North America (Cranoglanididae and Ictaluridae) were recovered in a closely related clade. The branch lengths separating these seven families were relatively short, suggesting their rapid diversification within these families over a relatively short period of time. To test this, we estimated the divergence time using MCMCTree.

The MCMCtree analysis revealed that the divergence of South American catfishes from their relatives included in this study occurred around 114 MYA, with a 95% confidence interval of 103 – 121 MYA (Fig. 2). This coincides with the early Cretaceous, when the complete separation of South America and Africa is believed to have occurred around 100 MYA (Reguero et al. 2021). Following that, between approximately 84 - 77 MYA, the divergence of other major families of catfishes of Africa and Asia occurred. Overall, the results suggest that the divergence of catfish species was mainly mediated by vicariance events, closely linked to the breakup of Gondwana, with Africa/Gondwana potentially being the origin of catfishes (also see supplementary figs. 2, 3 & 4 for divergence estimation with prior and convergence plots).

**Fig. 2:**
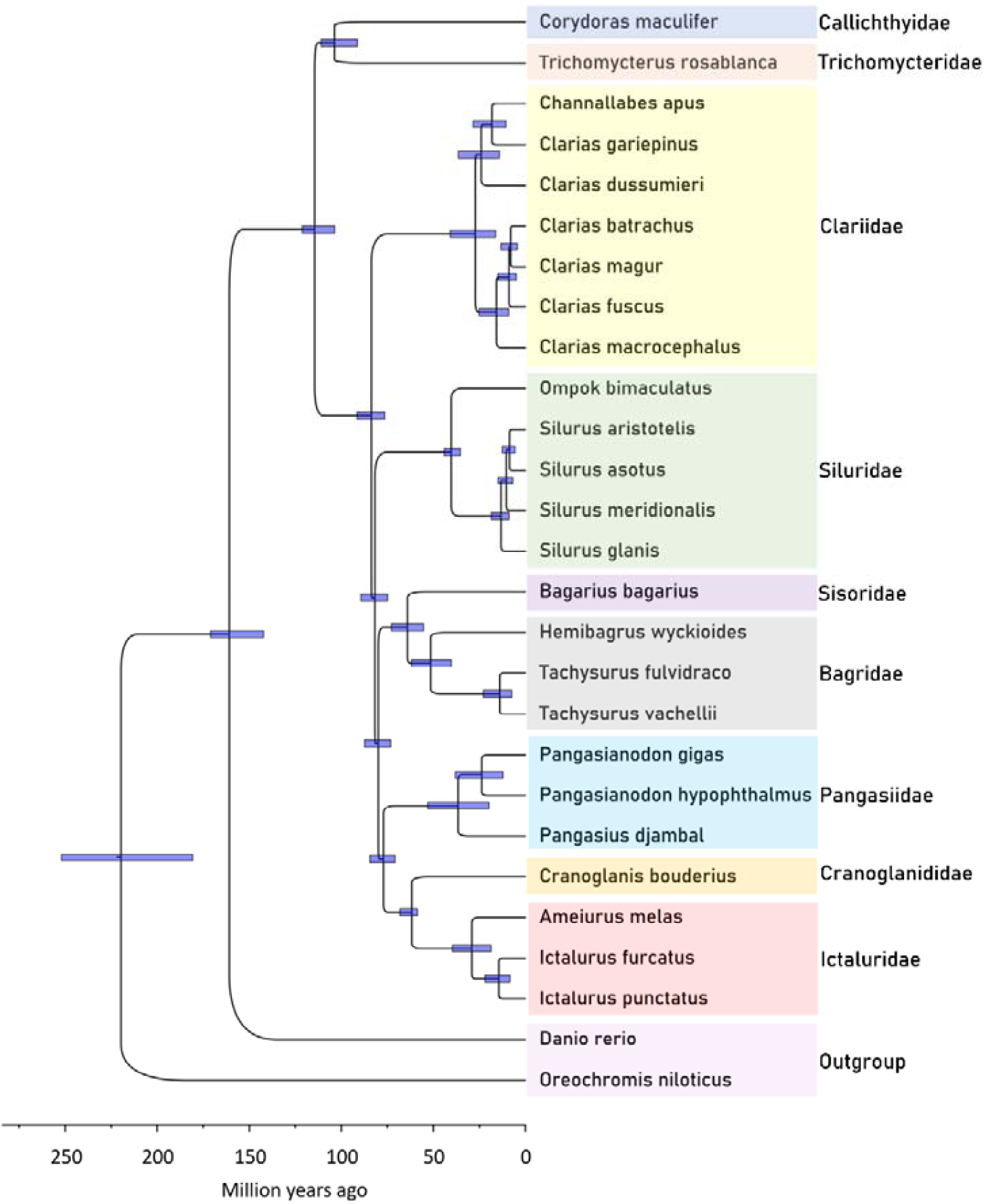
Species tree with divergence time estimation of catfish species belonging to nine families plus two outgroup species. The bars represent the 95% confidence intervals for the estimated divergence times.

### 3. Positive selection analysis

The comparison between five catfish families revealed that Clariidae has the greatest number of genes (121) under positive selection compared to the other four families, while Bagridae had the least (70) (Fig. 3b). Since there were more than one site with a probability of >95% positive selection in certain genes, we also counted the total number of such sites for each lineage and found similar results, where Clariidae had the highest number of positively selected sites in total (194 sites from 121 genes). In the other four families, most of the positively selected genes had only single positively selected site with shared mutation (BEB >95%) among the species of the respective family, while Clariidae had the greatest number of genes that had more than one positively selected site with shared mutations (Fig. 3c). Based on the functions of these genes determined through various genetic studies on model organisms, we grouped them based on the traits they are involved in and their interactions (Fig. 3a).

**Fig. 3:**
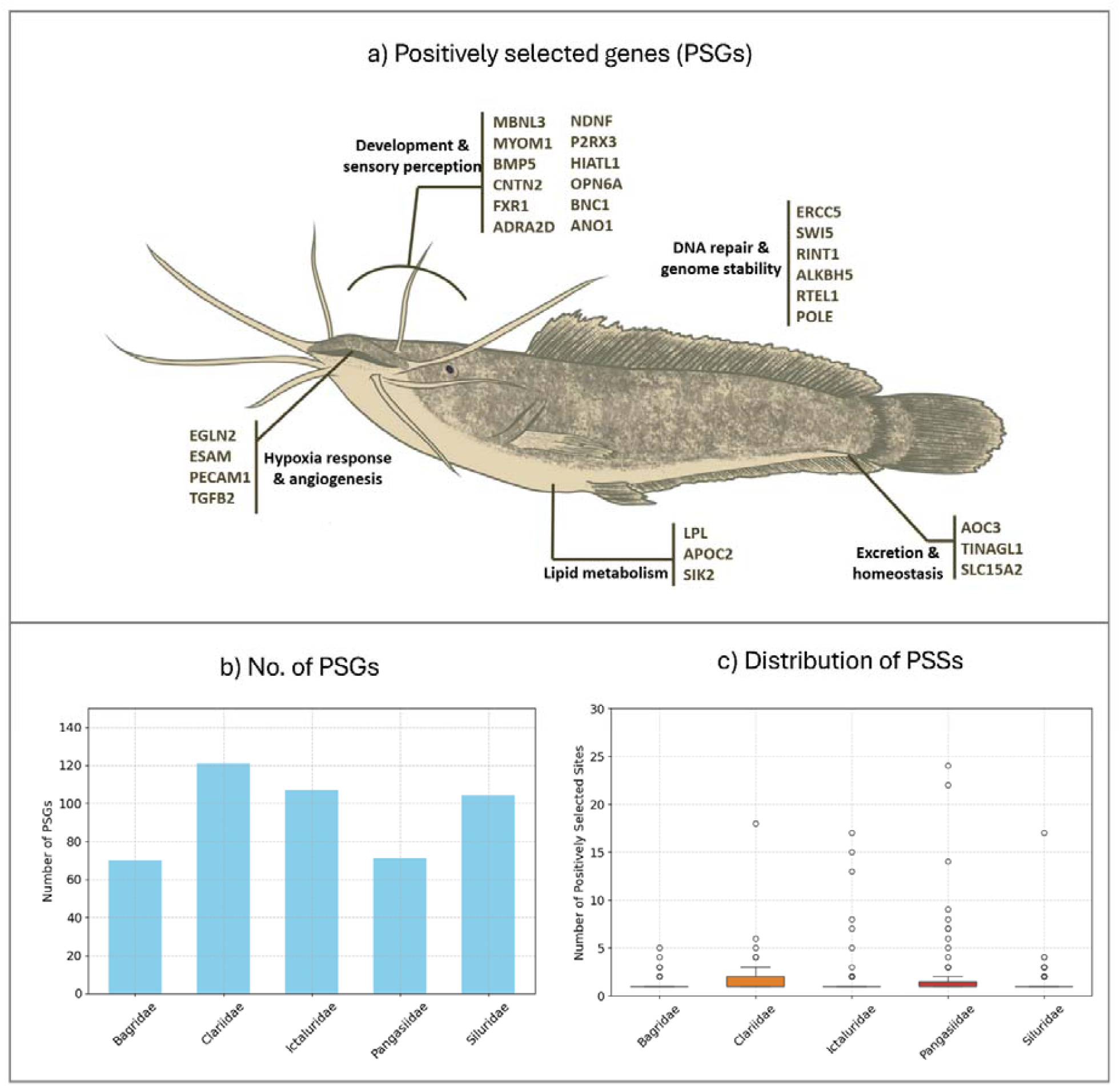
a) Major categories of positively selected genes in Clariids. b) No of positively selected genes across five families of catfishes. c) No. of positively selected sites for the PSGs in the five catfish families.

#### a) Adaptation to hypoxic conditions

In the transition from aquatic to terrestrial environments, a significant challenge for fishes is the efficient transport of oxygen to cells. While lungfishes have evolved true lungs, clariids lack such adaptations. Oxygen transport in the absence of lungs necessitates compensatory molecular mechanisms, including enhanced oxygen delivery and protection against hypoxic stress. Under normoxic conditions, HIF-1α undergoes proteasomal degradation, mediated by its hydroxylation via EGLN2, which promotes VHL-mediated ubiquitination (Semenza 2007) (Fig. 4a). In our study, we identified positive selection in the EGLN2 gene, with a shared mutation at 21^st^ position in clariids (Fig. 4b), suggesting its adaptive role in regulating HIF-1α activity (Fig. 4a). However, during hypoxia, HIF-1 drives the transcriptional activation of various genes involved in cellular growth, angiogenesis, erythropoiesis, and transcriptional regulation (Lee at al. 2004). These responses promote oxygen delivery to hypoxic regions. Genes such as Platelet Endothelial Cell Adhesion Molecule 1 (PECAM1), Endothelial Cell-Selective Adhesion Molecule (ESAM) and CPOX were also positively selected (Fig. 4b). PECAM1 plays a critical role in angiogenesis (Ferrero et al. 1995) and ESAM supports vascular integrity (Duong et al. 2020), while CPOX is involved in heme biosynthesis (Dailey and Meissner 2013).

**Fig. 4:**
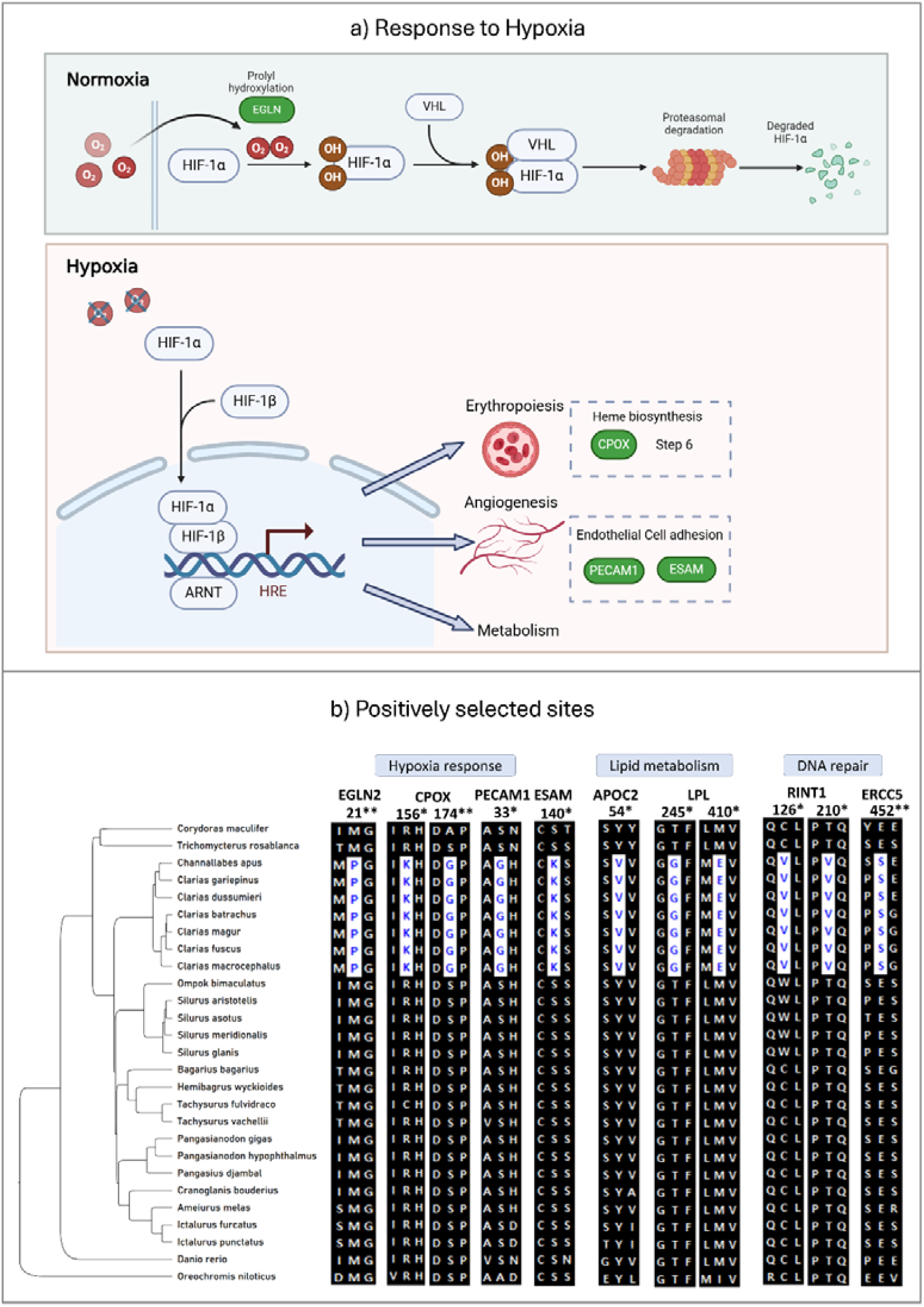
a) Function of HIF-1 alpha during normoxia and hypoxia, and positively selected genes involved in regulation of Hypoxia stress. The genes colored in green are positively selected in clariids. b) Shared mutations among clariids in significant PSGs of major categories. * indicates the posterior probability of the site being positively selected between 95% and 99%, ** indicates the posterior probability >99%.

#### b) Development and sensory perception

Seventeen genes directly involved in cellular and organ-level development processes, such as neurodevelopment, muscle development, regeneration, bone development, and reproduction, were positively selected in clariids. This includes genes related to muscle development include Muscleblind-like 3 (MBNL3), which supports skeletal muscle development and regeneration (Poulos et al. 2013), and Myomesin 1a (skelemin), crucial for striated muscle and heart development (Reddy et al. 2008) (supplementary fig. 5). Given that clariids often traverse land, move upstream against currents, and burrow in riverbanks, these genes likely enhance muscle strength and coordination. Additionally, Myomesin 1a may aid in heart development, as hypoxic conditions necessitate increased blood production and a stronger heart to support enhanced blood flow. TGFB2, which plays critical roles in developmental processes (Azhar et al. 2011), including valve remodelling during heart development was also found to be positively selected at multiple positions (110, 111, 120, 179 & 233) (Supplementary fig. 6). Similarly, positive selection in bone morphogenetic protein BMP5, essential for bone development (Ducy and Karsenty 2000), may reflect adaptations supporting their terrestrial and aquatic activities.

Five genes involved in neurodevelopment and associated functions were also positively selected. These include Contactin 2 (CNTN2) for axon connections (Mohebiany et al. 2013), Fragile X mental retardation (FXR1) for dendrite formation (Shen et al. 2021), Adrenergic, alpha-2D receptor (ADRA2D) (Kalapesi at al. 2005) for neuroprotection, and Neuron-derived neurotrophic factor (NDNF) for neuromuscular functions (Ohashi et al. 2014; Joki et al. 2015). Also, Purinergic receptor P2X, ligand-gated ion channel 3a (P2RX3), associated with pain reception (Jarvis et al. 2002), and Opsin 6 (group member a), a subclass of melanopsin, expressed in photoreceptors (Bertolesi et al. 2022) were positively selected (See Supplementary fig. 6 for details).

#### c) Cellular homeostasis

The transition from aquatic to terrestrial environments necessitates robust physiological mechanisms to maintain cellular homeostasis. These include regulating ionic and organic molecule concentrations, detoxifying harmful substances, and adapting excretion strategies to the constraints of a terrestrial habitat. Positive selection was observed in the AOC3 gene, encoding vascular adhesion protein (VAP-1), which catalyzes oxidative deamination of amines, producing aldehyde, ammonia, and H₂O₂ (Smith and Vainio 2002). Its activity has been linked to vascular health and responses to hypoxic stress (Zhang at al. 2017). Similarly, the TINAGL1 gene, expressed during kidney development, selectively regulates tubulogenesis, indicating its role in renal adaptation (Yoshioka at al. 2002). The SLC15A2 gene, part of the solute carrier family, mediates peptide reabsorption in the kidney, essential for maintaining metabolic balance under terrestrial conditions (Daniel and Kottra 2004). All these three genes demonstrated significant positive selection with shared mutations (Supplementary fig. 7).

#### d) Lipid metabolism

Lipoprotein lipase (LPL) and apolipoprotein C-II (ApoC-II) are critical genes in lipid metabolism, with LPL hydrolyzing triglycerides and very low-density lipoproteins (VLDL) to provide energy for tissues (Mead et al. 2002), while ApoC-II acts as its essential activator (Kinnunen et al. 1977). Studies have shown that fasting increases ApoC-II levels in mice (Rhee et al. 2006), and apoc2 mutant zebrafish exhibit dysregulated lipid metabolism and hyperlipidemia (Ka and Jin 2020). In this study both genes showed significant positive selection in clariids, with shared mutations at positions 54 (APOC2) and 245 & 410 (LPL) (Fig. 4b). Clariids are known to have higher fat content (Adebayo at al. 2016), and the positive selection of these genes likely aids in lipid storage and energy management, particularly under terrestrial conditions where energy reserves are critical for survival. Interestingly, SIK2, a gene hypothesized to regulate insulin sensitivity and glucose uptake, was also positively selected (Supplementary fig. 7). Its selection may act as a protective mechanism, potentially mitigating the adverse effects of increased fat storage, such as hyperlipidemia-induced insulin resistance (Sakamoto et al. 2018). Together, these results highlight the coordinated regulation of lipid metabolism and energy homeostasis in response to terrestrial environmental demands.

#### e) DNA repair

Transitioning to land exposes fish to environmental challenges such as UV radiation, which can lead to DNA damage. Additionally, terrestrial conditions often impose stressors like hypoxia, increasing intracellular oxidative stress and exacerbating DNA damage. In our study, six positively selected genes were identified, all associated with DNA repair, underscoring their significance in maintaining genomic stability under these conditions. We identified significant positive selection in Excision Repair Cross-Complementation Group 5 (ERCC5) gene (Fig. 4b), which is crucial for repairing UV-induced DNA damage. Mutations in this gene are known to cause xeroderma pigmentosum (Robbins 1988). A comparative study of surface-dwelling and cave-dwelling amphipods revealed relaxed selection and reduced expression of ERCC5 in cave-dwelling species, likely due to their lack of sunlight exposure (Carlini and Fong 2017). The positive selection of ERCC5 in clariids highlights its role in protecting against UV-induced DNA damage during terrestrial exposure.

Positive selection was also observed in other DNA repair-related genes. SWI5, a homolog involved in DNA recombination repair (Akamatsu et al. 2003), RAD50 Interactor 1 (RINT1), which regulates cell cycle control following DNA damage (Xiao et al. 2001), and ALKBH5, which repairs alkylation-induced DNA damage (Akula et al. 2021). Notably, ALKBH5 has been shown to be a direct transcriptional target of HIF-1α, linking it to cellular responses to hypoxia (Thalhammer et al. 2011). Additionally, the Regulator of Telomere Length 1 (RTEL1), known to maintain telomere integrity by resolving DNA secondary structures during repair (Vannier et al. 2014), and DNA Polymerase Epsilon (POLE), essential for DNA replication and repair, were also significantly positively selected (See Supplementary fig. 8 for details). Similar findings of positive selection in DNA repair genes by Li et al. (2018a) highlight their importance in genome stability and adaptation to novel conditions.

In addition to the above-mentioned categories, we observed significant positive selection in genes related to immunity and cellular processes as well, such as Complement component 8 (C8A), Raftlin family member 2 (RFTN2), Complement C2-like (C2L) and Prostaglandin E receptor 4 (PTGER4). While the functions of these genes are many and not limited to one, further investigation would reveal the complex interplay of multifunctional genes in the evolution of adaptive traits.

### 4. Gene Family Evolution analysis

Gene family evolution analysis using CAFE5 across 28,223 orthogroups showed that 779 gene families were significantly expanded or contracted (adjusted p-value < 0.05). Among these, 205 were expanded specifically in the family Clariidae, and 74 families were contracted. Functional annotation of these expanded families in clariids through eggNOG-mapper revealed key roles in oxygen transport, xenobiotic degradation, immunity, fertility, neurodevelopment, and essential cellular functions (Fig. 5a). Notably, Myoglobin, a critical gene for oxygen transport and storage, is typically found in single-copy in most vertebrates. However, in clariids, this gene underwent substantial expansion (Figure 5-a&b), with copy numbers ranging from 9 to 13 (Supplementary fig. 10). CAFE prediction indicates that this expansion probably occurred in the common ancestor of clariids, reaching approximately 10 copies.

**Fig. 5:**
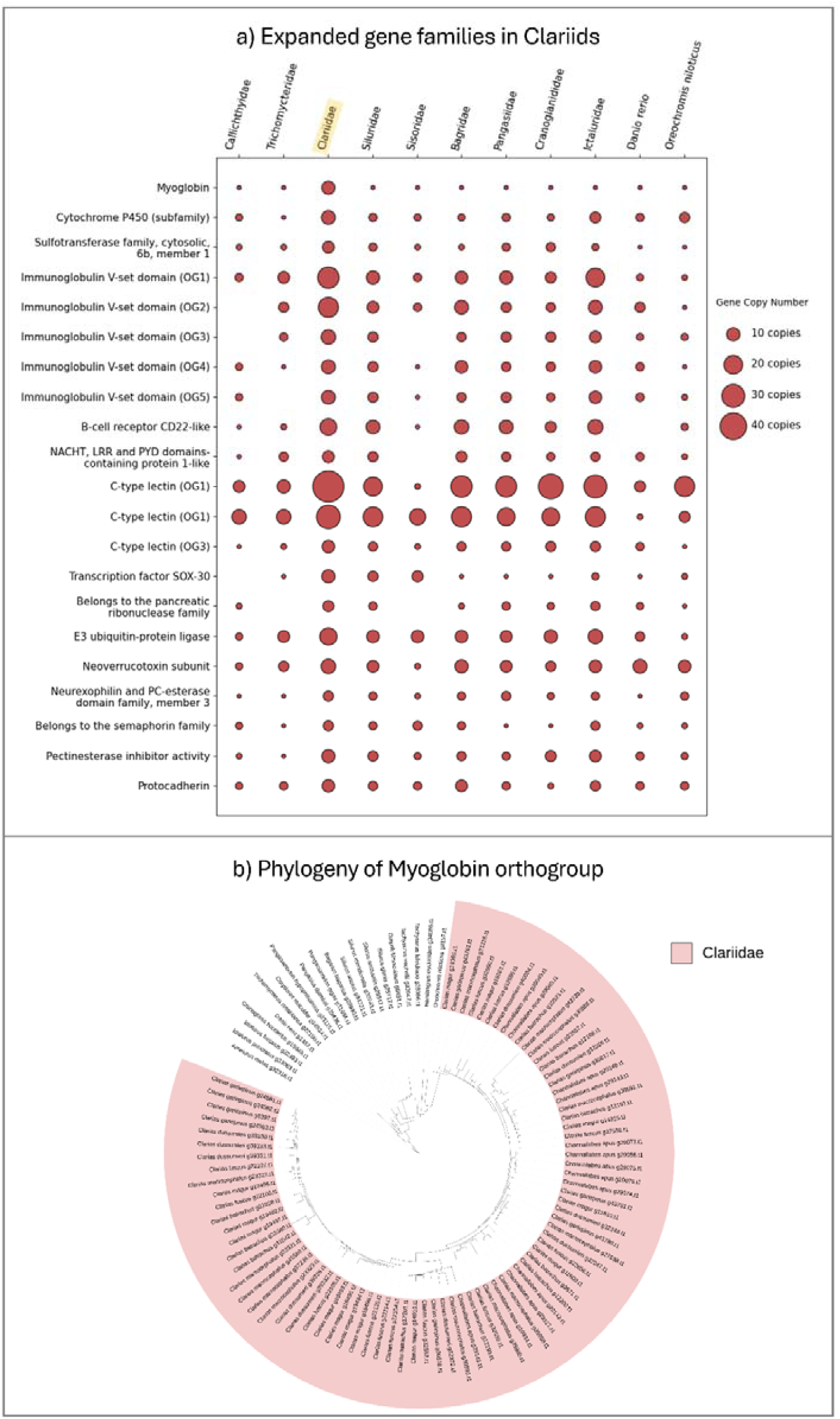
a) Expansion of major genes families in clariids relative to other catfish families and outgroups. b) Phylogeny of Myoglobin orthogroup depicting massive expansions in Clariids.

In addition, several orthogroups associated with immunity, such as Immunoglobulin V-set domain and C-type lectin, displayed significant expansions. Genes involved in xenobiotic degradation, including members of the Cytochrome P450 subfamily and the Sulfotransferase family, cytosolic, 6b, member 1, also expanded. These expansions likely highlight adaptations to terrestrial environments, where combating various pathogens and parasites on land, and effective xenobiotic processing is critical. Genes associated with crucial cellular processes, such as E3 ubiquitin-protein ligase, Semaphorin, Protocadherin, and Pectinesterase inhibitor, also exhibited notable expansions in clariids, further emphasizing the lineage’s unique evolutionary adaptations.

GO enrichment analysis revealed high levels of enrichment of several terms in the expanded gene families of Clariids. They were predominantly related to immunity such as GO:0002223 (stimulatory C-type lectin receptor signaling pathway), GO:0006958 (complement activation, classical pathway) and GO:0061760 (antifungal innate immune response), to name a few. Notably, the GO terms GO:0015671 (oxygen transport) and GO:1990785 (response to water-immersion restraint stress) were also enriched several folds in all the 7 clariid species (Supplementary fig.11). These findings align with the observed gene family expansions, substantiating the importance of these functions in adaptation of clariids to terrestrial life.

### Summary and concluding remarks

In this study, we explored the genomic signatures underlying ecological diversification and the transition to extreme or novel conditions in clariids. By examining how their genomes evolved in response to multiple stressors, we identified over 100 genes under strong positive selection with high statistical significance. These genes are linked to critical adaptive functions, including hypoxia response, development, sensory system, lipid metabolism, and DNA repair. Additionally, we uncovered rapidly expanding gene families, such as Myoglobin, immune-related genes, and xenobiotic degradation, highlighting their pivotal roles in facilitating adaptation to both aquatic and terrestrial environments.

Previous genomic studies on clariids have also identified signatures of terrestrial adaptation. For instance, Li et al. (2018b) highlighted the expansion of myoglobin genes and transcriptional upregulation of hemoglobin genes in *Clarias batrachus* compared to channel catfish (*Ictalurus punctatus*). Similarly, Kushwaha (2021) identified signatures for adaptations to hypoxia, ammonia tolerance, urea cycle regulation, vision, locomotion, and detoxification in *Clarias magur*. However, no studies have comprehensively investigated the genome evolution in clariids for terrestrialization. To address this, our study incorporated 25 catfish species, including seven clariids, with a more robust and meticulous phylogenomic approach. Unlike previous studies that relied on protein-coding sequences annotated with different bioinformatics pipelines by various scientific groups, we downloaded and re-annotated the genomes of all species using the same bioinformatics pipeline to minimize technical artifacts.

Certain genes have displayed signs of positive selection with higher dN/dS ratios and shared non-synonymous mutations in clariids, linked to traits like response to hypoxia (HIF-1α) and ionic regulation, but did not pass LRT test. For such genes, BEB analysis revealed multiple sites with probability of sites under selection exceeding 95% (Supplementary fig. 9), suggesting that specific residues might be under strong selection even if the overall gene-level signal is weaker. Further investigation of such genes could refine statistical models and enhance existing frameworks (Yang et al. 2005). Moreover, detailed analysis of orthologs and paralogs following gene duplication events could provide insights into evolutionary pressures and neofunctionalization. Additionally, several positively selected genes, including homeobox proteins and master regulators, highlight the complex pathways driving adaptation, though their precise roles are unclear.

Genome evolution during ecological diversification enables species to develop specialized traits for adapting to novel environments, granting them a competitive advantage. These evolutionary novelties, driven by selection pressures, also enhance the ability to adapt and invade rapidly to new habitats. *Clarias gariepinus* and *Clarias batrachus* are among the most invasive fish species, with *C. batrachus* listed among the world’s 100 worst invasive species (Global Invasive Species Database, 2025). Their adaptations to extreme conditions, such as surviving droughts and polluted waters, have facilitated their invasion of ecosystems and outcompeting native species, underscoring their status as “ideal weeds.” Large-scale comparative genomics of species transitioning to novel niches offers valuable insights into the mechanisms that enable such invasiveness. This area remains underexplored and could shed light on the evolutionary processes underlying rapid adaptation and ecological dominance in invasive species.

While genomic studies on other clariids align with our findings, these differences in gene family expansions, contractions, and positive selection likely stem from variations in bioinformatic pipelines and data quality. Future advancements in sequencing technologies, reduced costs, and more standardized computational workflows, including the generation of chromosome-level assemblies, will enhance accuracy and reproducibility. Such developments will significantly aid in understanding eco-evolutionary processes, conserving biodiversity, and advancing applications in agriculture and aquaculture.

Finally, this study shows how genomic adaptations enable air-breathing catfishes to thrive across extreme and novel environments, both in water and on land. Our findings will also help future exploration of genome evolution in other invasive and resilient species. As the field advances with improved methodologies and richer datasets, such studies will continue to illuminate the evolutionary mechanisms behind ecological divergence and their implications for biodiversity and ecosystem dynamics.

## Acknowledgements

The authors would like to thank Dr. Divya Tej Sowpati and Dr. Jahnavi Joshi for their very valuable insights. The authors also thank the Next generation sequencing facility of CSIR-Centre for Cellular and Molecular Biology, Hyderabad for their assistance in generating the sequence data, and Mr. Yashwant Singh Panwar for the catfish illustration.

## Data availability

All the raw sequences generated for this study have been submitted to NCBI SRA under BioProject ID PRJNA1219819. The Biosample IDs are SAMN46701642 for *Clarias gariepinus* and SAMN46701643 for *Clarias dussumieri*. The assembled genomes of *C. gariepinus* and *C. dussumieri* have been submitted to NCBI Genbank under accession numbers JBLMLW000000000 and JBLMLV000000000 respectively.

## Supplementary information

## Supplementary methods

### 1. DNA isolation and sequencing

To isolate DNA, fin clips and muscle tissue were collected from one individual each (sex unknown) of *Clarias gariepinus* and *Clarias dussumieri* from Kerala, India. Different protocols were used for DNA extraction based on the sequencing method. For long-read sequencing, DNA was extracted using the standard phenol-chloroform method. A total of 100 mg of tissue was finely minced and lysed for 4 hours at 56°C in 1 mL of lysis buffer (50 mM Tris-HCl, pH 8.0; 100 mM EDTA; 100 mM NaCl) containing 0.5% SDS and 20 mg/mL Proteinase K. The lysate was then centrifuged at 4,000g for 15 minutes, and the supernatant was transferred to a fresh Lobind tube. To the supernatant, an equal volume of phenol:chloroform:isoamyl alcohol (25:24:1) was added and incubated for 10 minutes on a rotating wheel, followed by centrifugation at 10,000g for 10 minutes. The aqueous phase was transferred to a fresh Lobind tube, and the previous step was repeated. The final aqueous phase was further purified by incubating it with an equal volume of chloroform:isoamyl alcohol (24:1) for 10 minutes, followed by centrifugation at 10,000g for 10 minutes. The supernatant was then transferred to a 1.5 mL centrifuge tube and mixed gently with 0.05 volume of 5M NaCl and 0.7 volume of isopropanol. The mixture was incubated at-20°C for 8 hours and centrifuged at 10,000g for 30 minutes. The resulting DNA pellet was washed twice, first with 1 mL of 70% ethanol and then with 0.5 mL of 100% ethanol. After air-drying to remove any residual ethanol, the pellet was dissolved in 1X TE buffer at 56°C. For short-read sequencing, DNA was isolated using Qiagen’s Blood and Tissue DNA isolation kit, following the manufacturer’s protocol.

Whole-genome sequencing was performed at the Next Generation Sequencing facility of CSIR-Centre for Cellular and Molecular Biology, India. For long-read sequencing, libraries were prepared using Oxford Nanopore Technology’s Ligation Sequencing Kit V14 (SQK-LSK114), following the manufacturer’s protocol, and sequencing was carried out on a PromethION device. The sequencing reads were basecalled using the Guppy basecaller. For short-read sequencing, libraries were prepared using Illumina’s TruSeq DNA PCR-Free kit, and sequencing was performed on a NovaSeq 6000 device.

### 2. Genome assembly

Initial assembly was performed using only the long-read data. First, we removed the reads whose Phred quality score was below 9. We then assembled the genome using using Canu v2.2 (Koren et al. 2017) with the parameters used were {genomeSize=1g rawErrorRate=0.30 correctedErrorRate=0.12 corMinCoverage=4 corMhapFilterThreshold=0.0000000002 corMhapOptions=“--threshold 0.80--num-hashes 512--num-min-matches 3--ordered-sketch-size 1000--ordered-kmer-size 14--min-olap-length 2000--repeat-idf-scale 50” mhapMemory=192g mhapBlockSize=500 ovlMerDistinct=0.975 useGrid=false}. Second, the raw short reads were quality trimmed using fastp v0.23.2 (Chen et al. 2023) with the parameters {--detect_adapter_for_pe--trim_poly_g-5-3-l 51}. The trimmed short reads were supplied to polish the primary assembly using Polca pipeline of Masurca (Zimin et al. 2013). The polished genome was quality checked first with Quast v5.2.0 (Mikheenko et al. 2018) for the assembly stats, and then for completeness using BUSCO v5.7.1 (Manni et al. 2021), with Actinopterygii_odb10 dataset. The assembly summary was visualized using blobtoolkit v4.4.0 (Challis et al. 2020).

### 3. Repeat masking

From the repeat masking step, we followed the same pipeline till functional annotation for all the 27 species included in this study. We first created a custom repeat library for each species using Repeatmodeler v2.0.5 (Flynn et al. 2020). We then masked the genomes using RepeatMasker v4.1.4 (Smit et al. 2013) with the parameters {-nolow-s-xsmall-e rmblast} and with LTRStruct pipeline. The version of the search engine used was RMBLAST 2.13.0+.

### 4. Gene prediction

The soft-masked genome assemblies were used for gene prediction with GALBA v1.0.11 (Bruna et al. 2023), a fully automated gene prediction pipeline that employs AUGUSTUS in the absence of RNA-seq data. To train AUGUSTUS (Hoffe and Stanke 2019) with Miniprot (Li et al. 2023), we downloaded approximately 2 million protein sequences from the order Siluriformes from UniProt. For *D. rerio*, we used ∼111,000 protein sequences from Cypriniformes, and for *O. niloticus*, we used ∼385,000 sequences from Cichlidae. Gene predictions were filtered to remove unsupported predictions using the script selectSupportedSubsets.py. The remaining predictions were further refined by retaining only the longest isoforms using printLongestIsoforms.py. The final filtered gene predictions were processed with getAnnoFastaFromJoingenes.py (part of the GALBA pipeline) to extract coding sequences and translated protein sequences. The scripts selectSupportedSubsets.py and printLongestIsoforms.py were obtained from https://github.com/GaiusAugustus/BRAKER/tree/report/scripts/predictionAnalysis and were originally part of the BRAKER gene prediction pipeline. Finally, gene prediction completeness was assessed using BUSCO v5.7.1 with the *Actinopterygii_odb10* dataset.

### 5. Functional annotation

Functional annotation of the gene predictions was performed using eggNOG-mapper (emapper v2.1.12) (Cantalapiedra et al. 2021), based on eggNOG orthology data (eggNOG 5.0) (Huerta-Cepas et al. 2019). Sequence searches were conducted using DIAMOND v2.1.9 (Buchfink et al. 2015). To comprehensively obtain Gene Ontology (GO) terms for the predicted genes, we used FANTASIA (Martínez-Redondo et al. 2024), which annotates GO terms in protein sequence files using GOPredSim (Littmann et al. 2021) with the protein language model ProtT5.

### 6. Phylogenetic tree construction

Orthology assignment was performed using OrthoFinder v2.5.5 (Emms and Kelly 2019) to identify single-copy (SCO) and multi-copy orthologous sequences. SCOs were first aligned at the amino acid level using MAFFT v7.505 (Katoh and Stanley 2013). The corresponding nucleotide sequences were then extracted from the coding sequence files generated by GALBA and aligned separately using the codon-aware aligner MACSE v2.07 (Ranwez et al. 2018). The resulting alignments were trimmed with TrimAl v1.4.1 (Capella-Gutiérrez et al. 2009) to remove gap-rich regions. To construct a species tree, the trimmed alignments were concatenated into a supergene alignment using AMAS (Borowiec 2016), which was also used to generate a partition file. The concatenated alignment and partition file were then used for phylogenetic inference in IQ-TREE v2.3.2 (Minh et al. 2020; Hoang et al. 2018), with the parameters-alrt 1000-B 1000-bnni, allowing ModelFinder (Kalyaanamoorthy et al. 2017) to determine the best model for tree construction. The resulting species tree was visualized using FigTree v1.4.4.

### 7. Divergence time estimation

For species divergence estimation, we used the MCMCTree module of the PAML package v4.10.7 (Yang 2007). First, we extracted fourfold degenerate third codon positions (4DTv regions) from the concatenated supergene alignment. We then conducted approximate likelihood analyses following the MCMCTree tutorial (https://github.com/sabifo4/Tutorial_MCMCtree) (Álvarez-Carretero et al. 2022), performing the following steps: a) Rooted the species tree constructed with IQ-TREE and applied calibrations at seven nodes; six calibrations from TimeTree database (Kumar et al. 2022) and one fossil calibration point for Ictaluridae as used by Near et al. (2012). b) Calculated the substitution rate using the script calculate_rateprior.R. c) Obtained branch lengths, the gradient, and the Hessian, d) Ran MCMCTree by sampling from both the prior and posterior with multiple chains. e) Diagnosed the outputs for consistency. To ensure robustness, we performed six independent chains for both prior and posterior analyses. For each chain, we discarded the first 100,000 iterations as burn-in, then sampled 1,000 iterations per cycle until 20,000 samples were gathered, totaling 20 million iterations. Posterior estimation was conducted under the autocorrelated (GBM) rate model for each chain, and convergence between prior and posterior was assessed. Finally, we visualized the time-calibrated tree using FigTree v1.4.4.

### 8. Positive selection analysis

To identify genes under positive selection, we used the CODEML module of the PAML package v4.10.7. First, we obtained the trimmed alignments of single-copy orthologous sequences from Step 6. The Clariidae lineage was designated as the foreground, while all other lineages served as the background. CODEML was run under the branch-site model for each alignment separately, followed by a second run using the null model with a fixed omega value (fix_omega = 1, omega = 1). A likelihood ratio test (LRT) was performed by comparing the log-likelihood values between the null and alternative models to assess statistical significance for positive selection. Positively selected genes (PSGs) were filtered by having p-value <0.05 from LRT test, and requiring at least one site with a posterior probability > 0.95, as identified through Bayes Empirical Bayes (BEB) analysis. Gene functions were annotated using results from eggNOG-mapper. For comparison, the same analysis was conducted on four additional lineages: Bagridae, Ictaluridae, Pangasidae, and Siluridae. Lineages consisting of only a single species were not analyzed as foreground branches due to limited comparative power but were always included in the background set in all analyses.

### 9. Gene family evolution analysis

We used CAFE5 (Mendes et al. 2020) to analyze multi-copy gene families that have significantly expanded or contracted in Clariidae. The gene count file from OrthoFinder results and the ultrametric tree obtained from MCMCTree analysis were used as input for CAFE. Gene count data included 28,199 families after omitting 24 families with the largest size differentials between the minimum and maximum counts. First, we filtered out gene families that had evolved significantly (adjusted p-value < 0.05, Benjamini & Hochberg’s method). We then further refined the dataset by selecting orthogroups that exhibited expansions specifically in Clariidae. The functions of these expanded orthogroups were annotated using results from eggNOG-mapper.

### 10. Gene ontology enrichment analysis

To identify Gene Ontology (GO) terms enriched in the expanded gene families of Clariids, we conducted GO enrichment analysis using the TopGO R package (Alexa Rahnenführer 2009). The entire GO annotations of the respective clariid species were retrieved using FANTASIA, and were used as reference set. The significantly overrepresented GO terms were identified with adjusted P value < 0.05 (Benjamini & Hochberg’s method). The enrichment score was calculated by dividing the number of annotations in the expanded gene families by the expected number of annotations.

---

**Supplementary Fig. 1:**
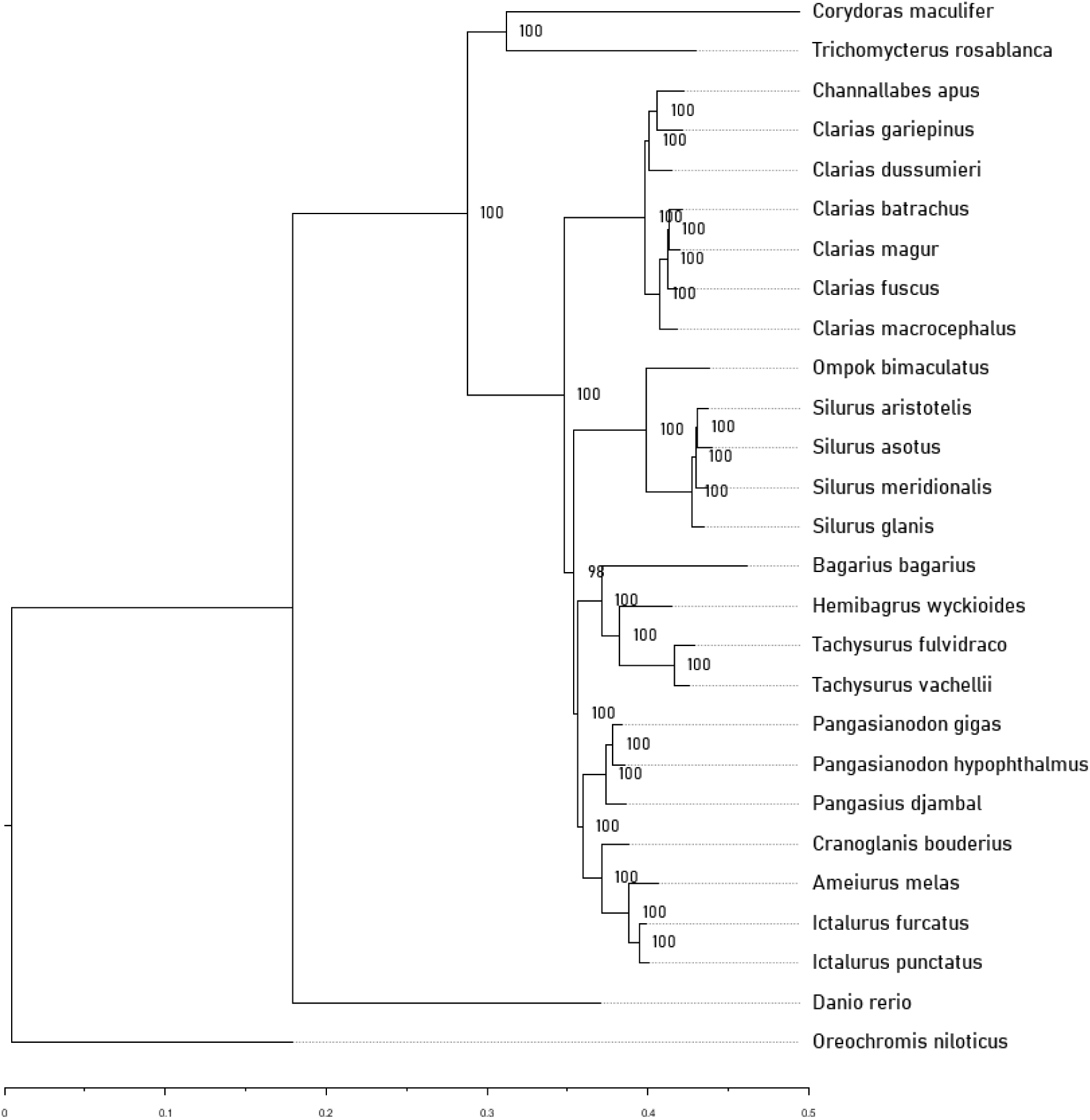
Phylogenetic tree constructed using IQTree. The node labels indicate the bootstrap support values.

**Supplementary Fig. 2:**
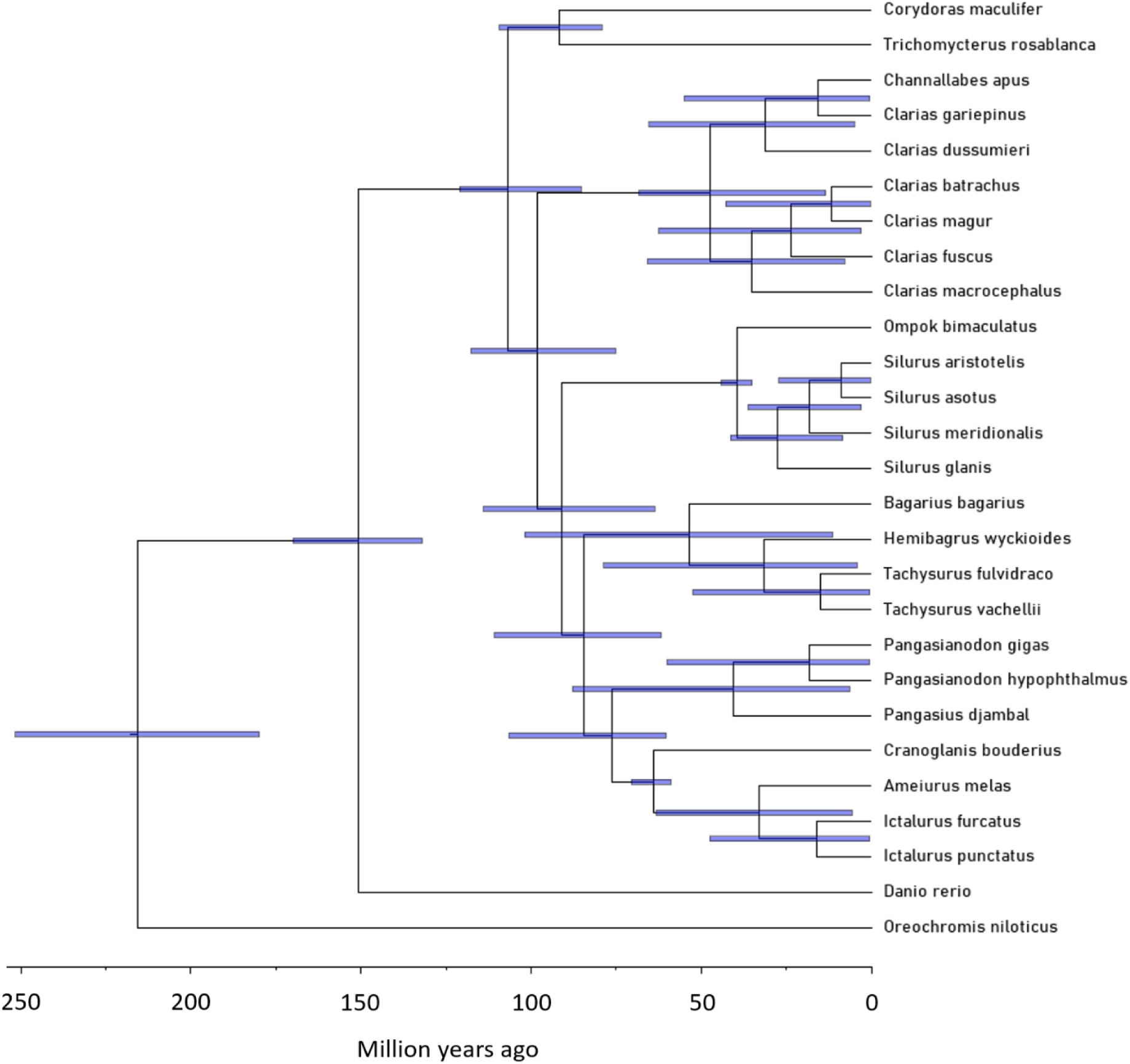
Divergence time estimation using Prior (without sequence data) using MCMCTree. The bars represent the 95% confidence intervals for the estimated divergence times.

**Supplementary Fig. 3:**
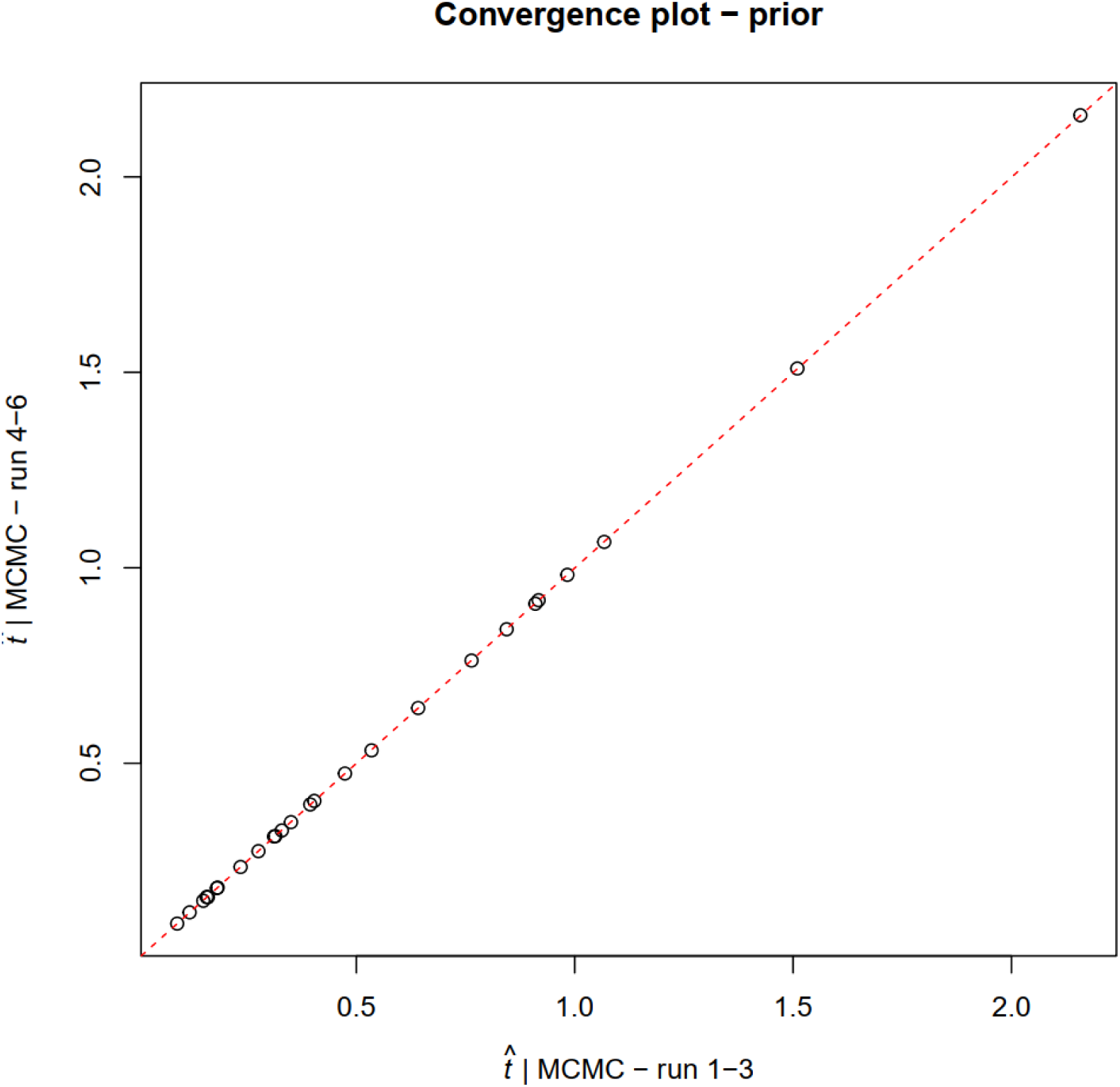
Convergence plot for the Prior divergence estimation of 6 independent chains.

**Supplementary Fig. 4:**
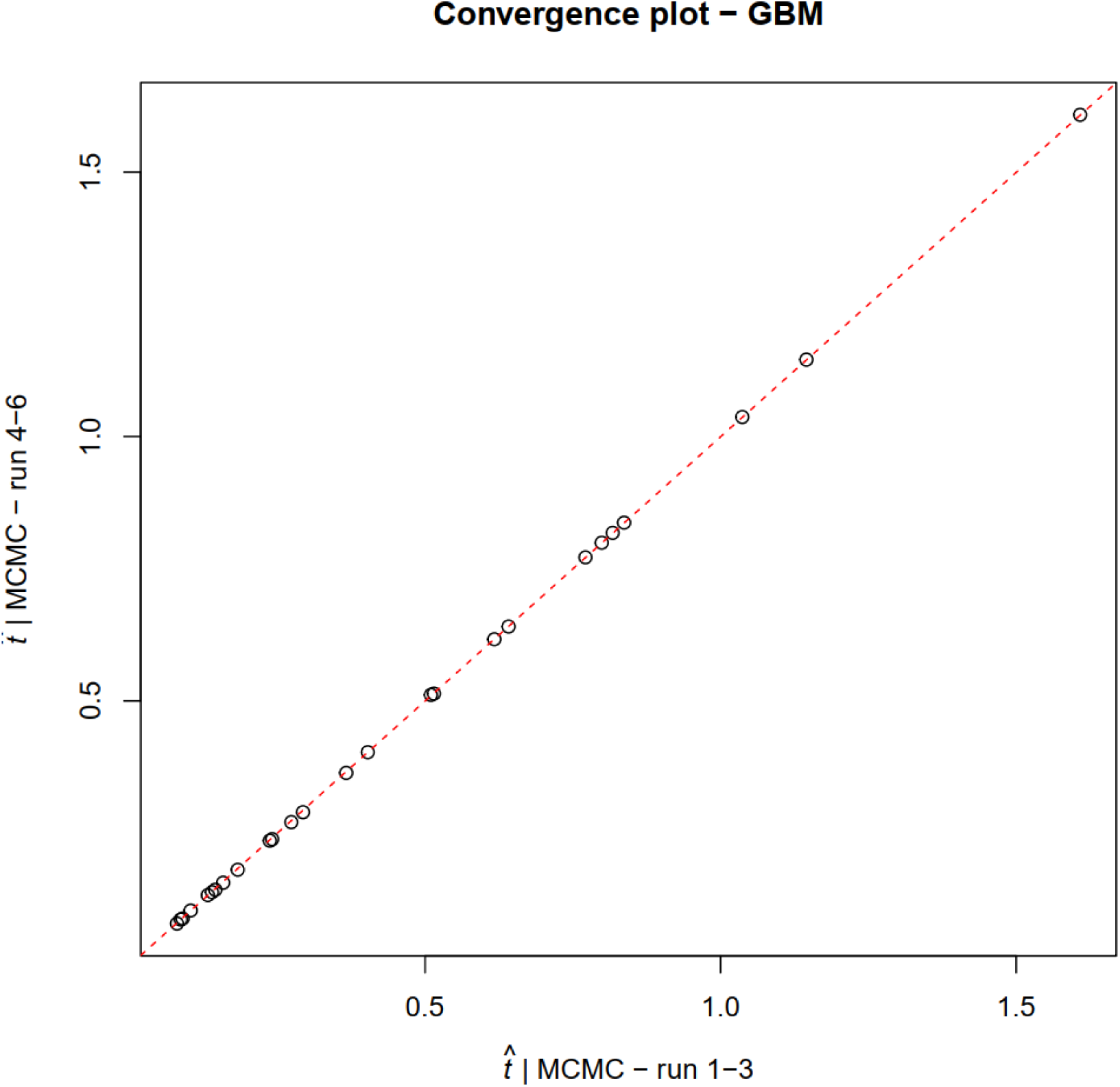
Convergence plot for the Posterior divergence estimation of 6 independent chains.

**Supplementary Fig. 5:**
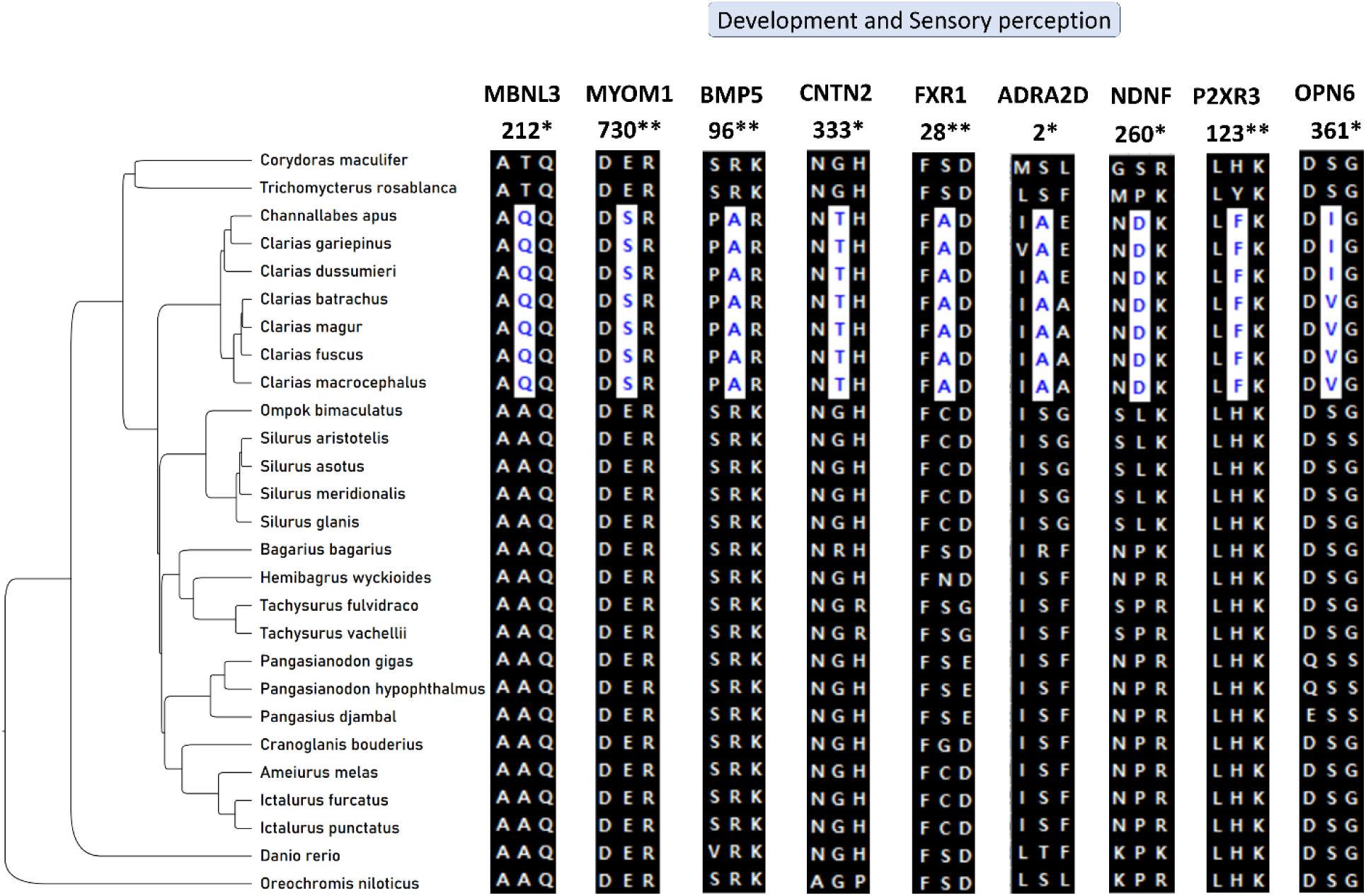
Shared mutations among clariids in significant PSGs involved in development and sensory perception. * indicates the posterior probability of the site being positively selected between 95% and 99%, ** indicates the posterior probability >99%.

**Supplementary Fig. 6:**
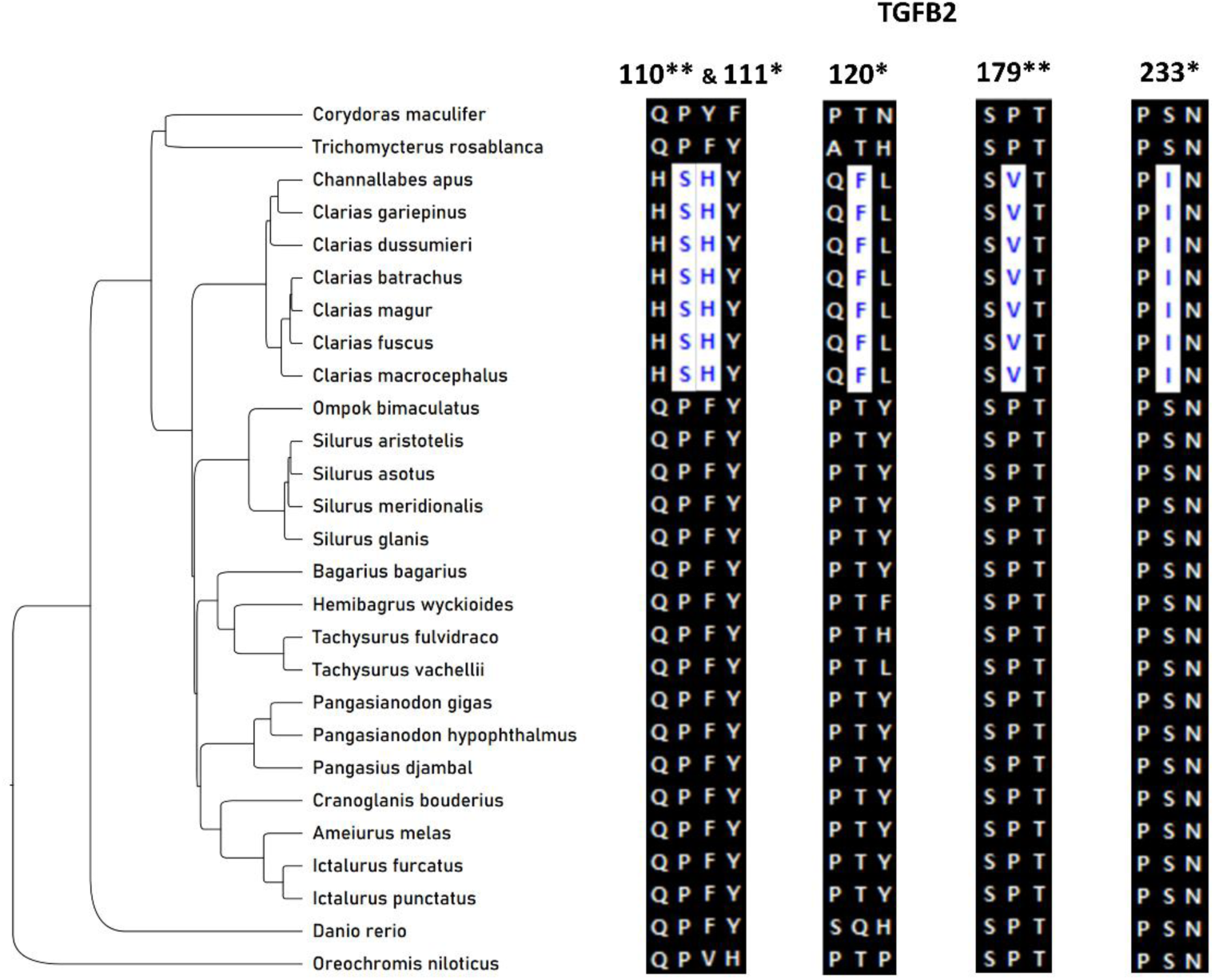
Shared mutations among clariids in TGFB2 gene. * indicates the posterior probability of the site being positively selected between 95% and 99%, ** indicates the posterior probability >99%.

**Supplementary Fig. 7:**
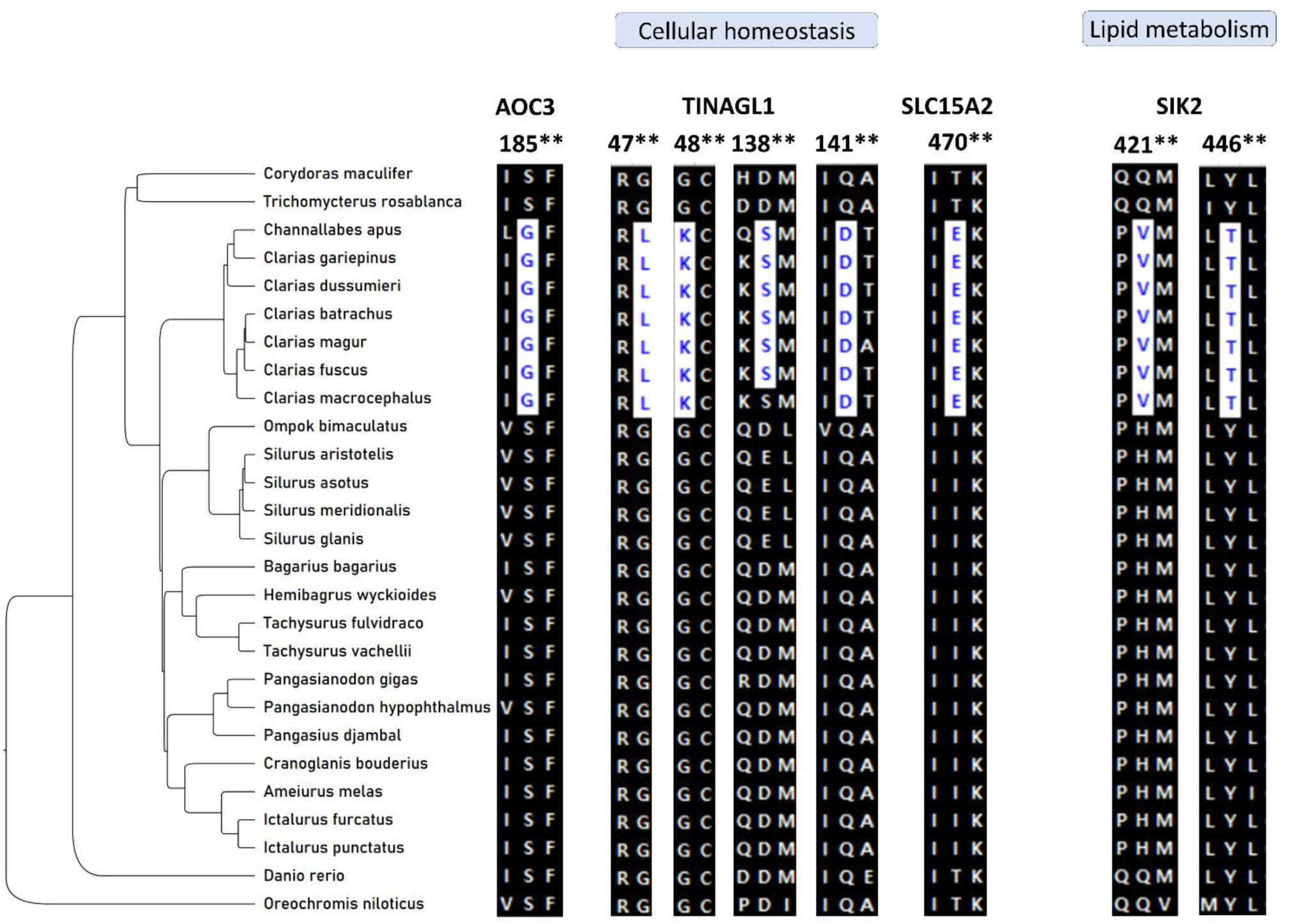
Shared mutations among clariids in significant PSGs involved in cellular homeostasis and lipid metabolism. * indicates the posterior probability of the site being positively selected between 95% and 99%, ** indicates the posterior probability >99%.

**Supplementary Fig. 8:**
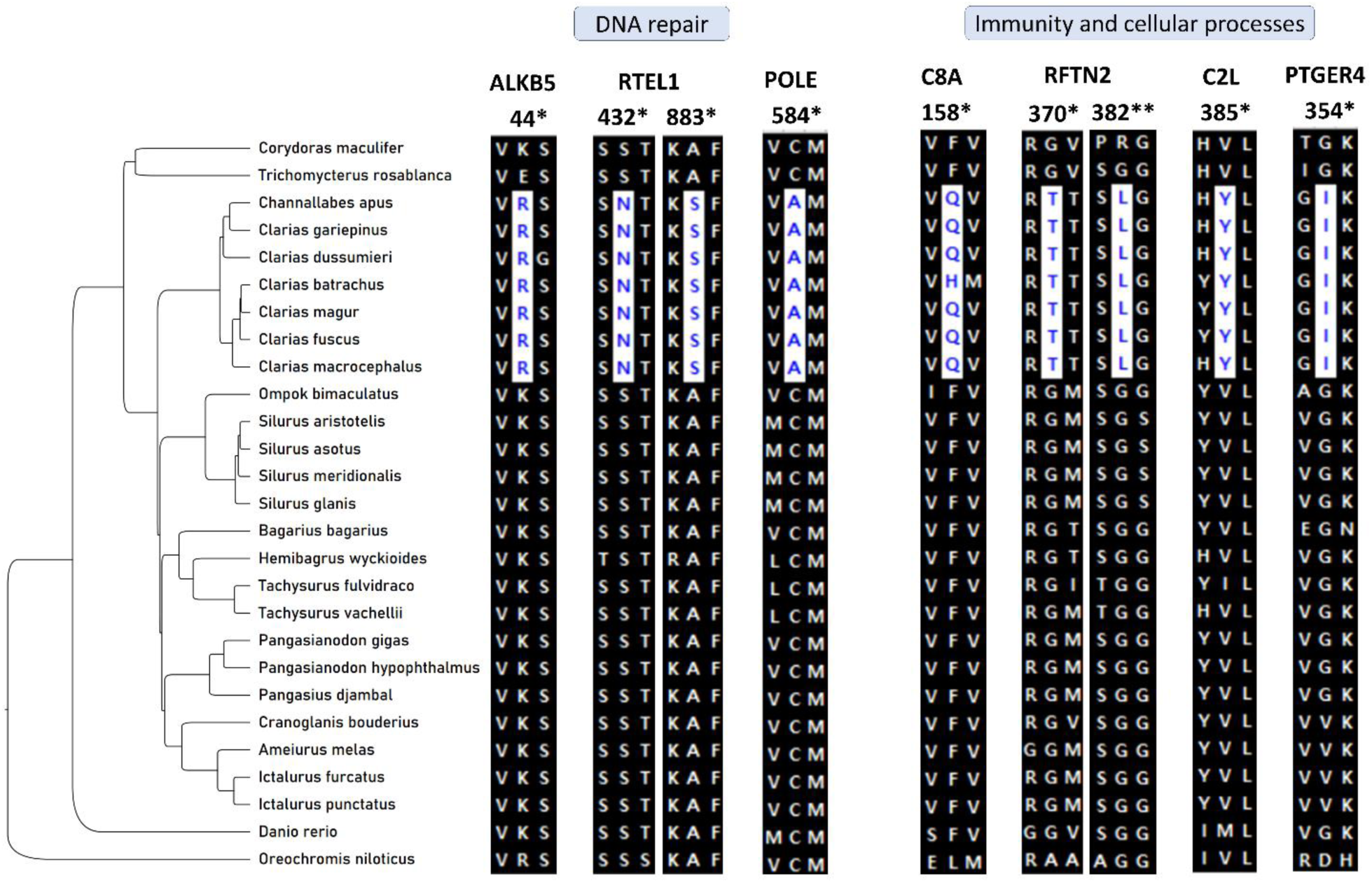
Shared mutations among clariids in significant PSGs involved in DNA repair and cellular processes. * indicates the posterior probability of the site being positively selected between 95% and 99%, ** indicates the posterior probability >99%.

**Supplementary Fig. 8:**
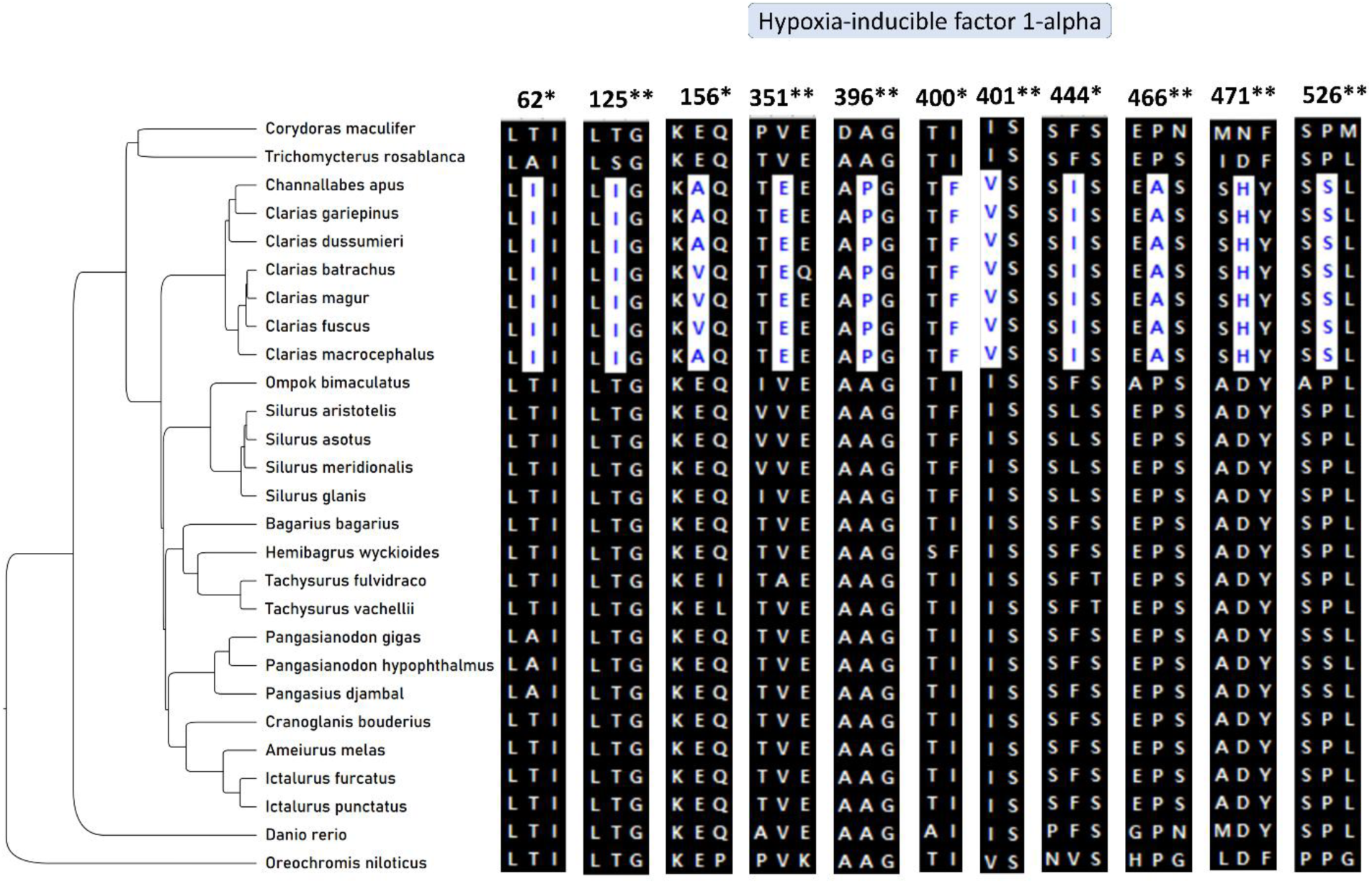
Shared mutations among clariids in HIF 1-alpha gene. * indicates the posterior probability of the site being positively selected between 95% and 99%, ** indicates the posterior probability >99%.

**Supplementary Fig. 10:**
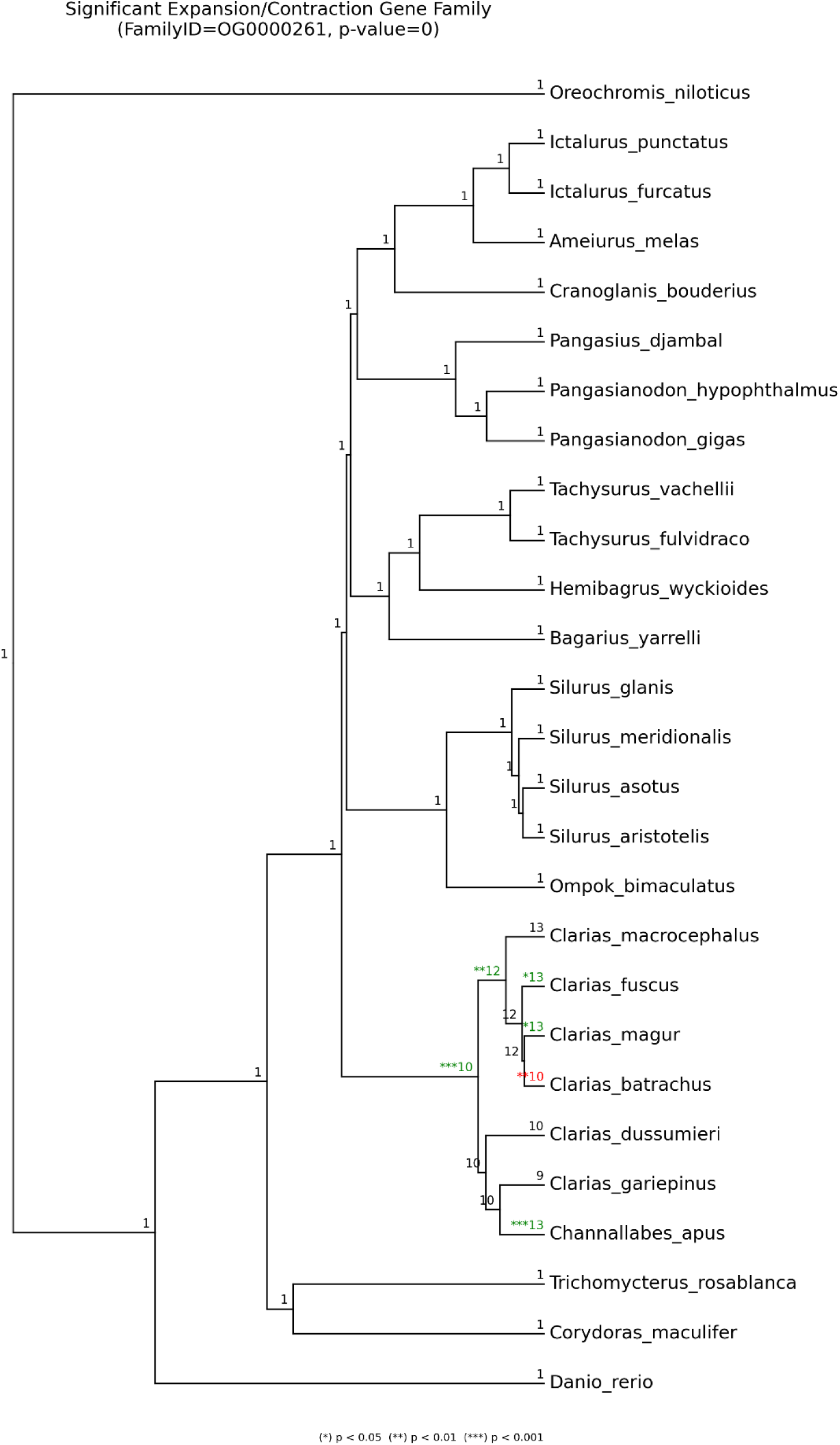
Expansion in the Myoglobin gene in Clariids. Green colour indicates expansion and red indicates contraction.

**Supplementary Fig. 11:**
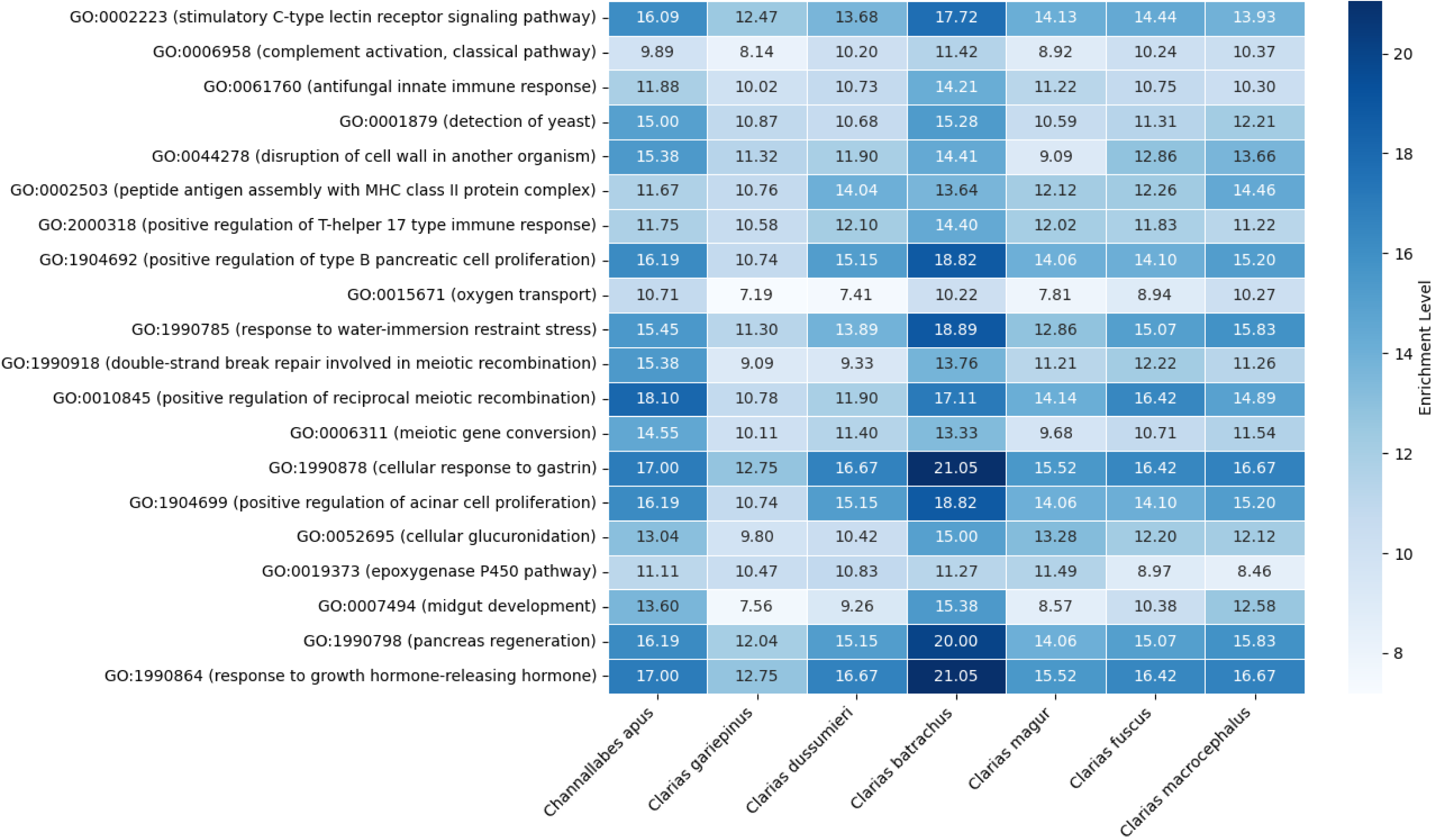
Enrichment levels of selected Gene Ontology terms in expanded gene families of clariid species.

**Supplementary Table 1:**
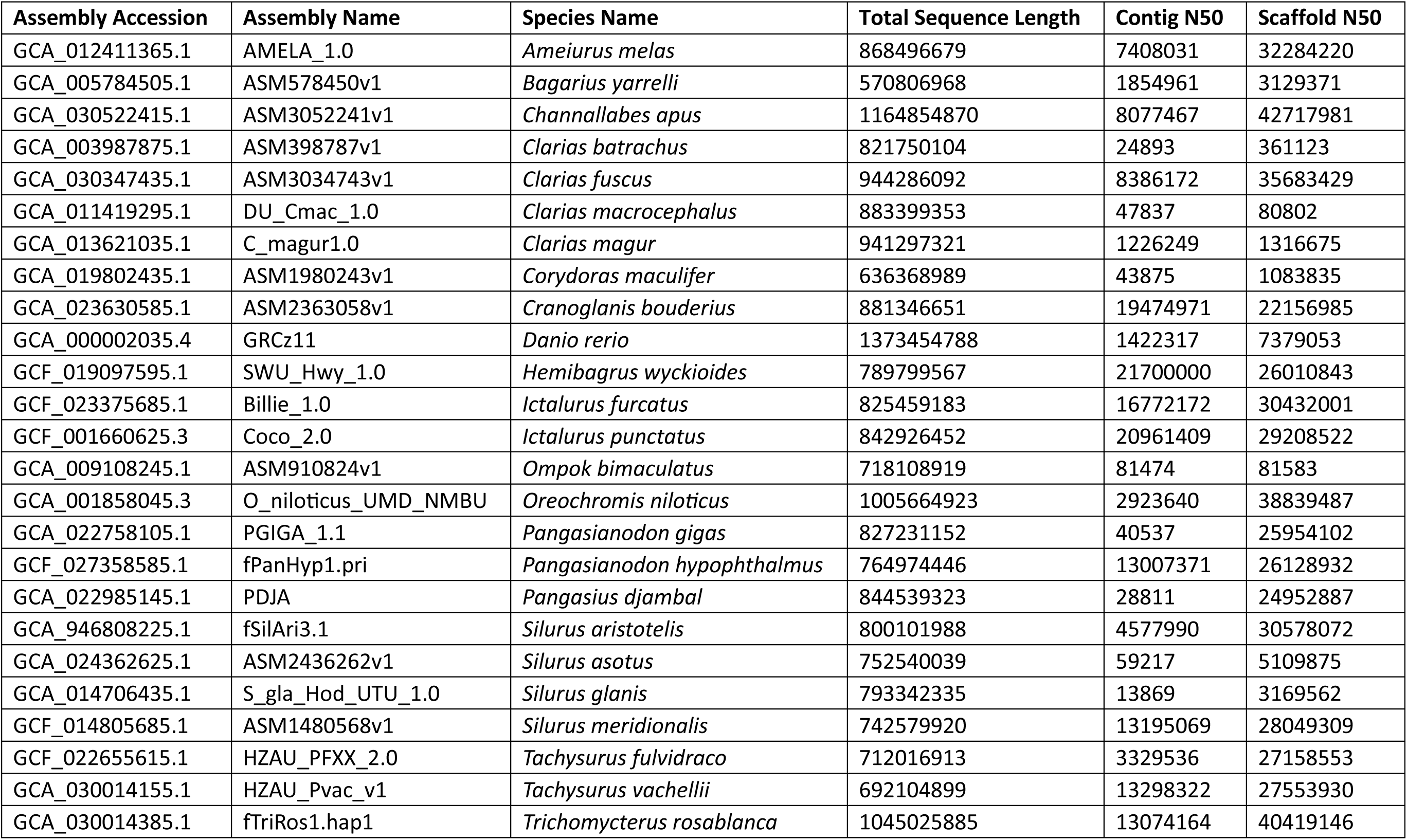
Summary of statistics of published genomes used in this study.

**Supplementary Table 2:** Repeat content characterization.

| **Species** | **DNA transposons** | **LTR** | **LINE** | **SINE** | **Unclassified** | **Others** | **Total** |
| --- | --- | --- | --- | --- | --- | --- | --- |
| *Ameiurus melas* | 17.35 | 7.21 | 4.9 | 0.59 | 12.92 | 1.85 | 44.82 |
| *Bagarius bagarius* | 8.89 | 2.85 | 4.25 | 3.36 | 9.08 | 1.24 | 29.67 |
| *Channallabes apus* | 17.45 | 13.66 | 9.05 | 0.69 | 12.33 | 1.04 | 54.22 |
| *Clarias batrachus* | 7.5 | 1.38 | 2.42 | 0 | 20.74 | 0.16 | 32.2 |
| *Clarias dussumieri* | 13.99 | 10.02 | 4.29 | 0.84 | 14.54 | 1.48 | 45.16 |
| *Clarias fuscus* | 14.17 | 7.45 | 5.69 | 0.51 | 11.54 | 1.46 | 40.82 |
| *Clarias gariepinus* | 13.89 | 8.99 | 10.35 | 0.38 | 12.37 | 1.25 | 47.23 |
| *Clarias macrocephalus* | 12.64 | 6.21 | 5.34 | 0.49 | 12.92 | 1.33 | 38.93 |
| *Clarias magur* | 12.47 | 8.53 | 6.12 | 0.61 | 13.54 | 1.47 | 42.74 |
| *Corydoras maculifer* | 4.89 | 8.88 | 8.71 | 1.09 | 7.54 | 0.41 | 31.52 |
| *Cranoglanis boudierius* | 21.87 | 5.68 | 4.67 | 0.6 | 14.1 | 1.04 | 47.96 |
| *Danio rerio* | 38.56 | 8.56 | 4.52 | 1.15 | 3.34 | 2.27 | 58.55 |
| *Hemibagrus wyckioides* | 10.94 | 2.86 | 4.07 | 0.46 | 10.99 | 5.81 | 35.13 |
| *Ictalurus furcatus* | 16.6 | 5.64 | 4.04 | 0.65 | 13.61 | 1.56 | 42.1 |
| *Ictalurus punctatus* | 17.54 | 5.91 | 4.11 | 0.63 | 14.03 | 1.67 | 43.89 |
| *Ompok bimaculatus* | 9.96 | 3.57 | 4.43 | 0.06 | 12.42 | 1.72 | 32.16 |
| *Oreochromis niloticus* | 10.92 | 8.56 | 8.22 | 0.26 | 11.38 | 0.96 | 40.27 |
| *Pangasianodon gigas* | 12.52 | 3.45 | 4.05 | 0.36 | 18.13 | 1.83 | 40.34 |
| *Pangasianodon hypophthalmus* | 10.74 | 3.32 | 3.76 | 0.5 | 16.86 | 1.62 | 36.8 |
| *Pangasius djambal* | 11.85 | 3.37 | 5.35 | 0.43 | 19.27 | 1.37 | 41.64 |
| *Silurus aristotelis* | 15.17 | 3.89 | 8.95 | 1.02 | 10.2 | 2.64 | 41.87 |
| *Silurus asotus* | 12.72 | 3.13 | 5.64 | 0.79 | 10.91 | 1.36 | 34.55 |
| *Silurus glanis* | 13.01 | 5.68 | 5.55 | 0.98 | 7.98 | 0.71 | 33.91 |
| *Silurus meridionalis* | 14.22 | 3.27 | 8.43 | 1.09 | 10.43 | 1.06 | 38.5 |
| *Tachysurus fulvidraco* | 8.27 | 6.47 | 7.06 | 0.35 | 10.76 | 2.23 | 35.14 |
| *Tachysurus vachellii* | 7.97 | 5.73 | 5.77 | 0.45 | 10.76 | 2.58 | 33.26 |
| *Trichomycterus rosablanca* | 22.21 | 8.61 | 4.46 | 1.94 | 16.3 | 1.51 | 55.03 |

**Supplementary Table 3:** Annotation statistics.

| Species | Genome size (Gb) | Total repeat content (%) | Gene count | Gene BUSCO completeness (%) |  |  |  |  |
| --- | --- | --- | --- | --- | --- | --- | --- | --- |
|  |  |  |  | Completeness | Single | Duplicated | Fragmented | Missing |
| *Ameiurus melas* | 0.87 | 44.82 | 27,701 | 96.1 | 94.8 | 1.3 | 1.3 | 2.6 |
| *Bagarius yarrelli* | 0.57 | 29.67 | 22,741 | 90.4 | 88.7 | 1.7 | 2.7 | 6.9 |
| *Channallabes apus* | 1.16 | 54.22 | 24,029 | 95.9 | 94.4 | 1.5 | 0.7 | 3.4 |
| *Clarias batrachus* | 0.82 | 32.2 | 24,981 | 88.3 | 86.6 | 1.7 | 3.7 | 8 |
| *Clarias dussumieri* | 1.08 | 45.16 | 26,023 | 96.2 | 93 | 3.2 | 0.6 | 3.2 |
| *Clarias fuscus* | 0.94 | 40.82 | 25,092 | 95.7 | 94.1 | 1.6 | 1 | 3.3 |
| *Clarias gariepinus* | 1.18 | 47.23 | 26,787 | 96.4 | 92 | 4.4 | 0.8 | 2.8 |
| *Clarias macrocephalus* | 0.88 | 38.93 | 26,589 | 89.5 | 87.7 | 1.8 | 4.9 | 5.6 |
| *Clarias magur* | 0.94 | 42.74 | 26,326 | 94.8 | 92.8 | 2 | 1.2 | 4 |
| *Corydoras maculifer* | 0.64 | 31.52 | 19,629 | 86.7 | 85.3 | 1.4 | 3.6 | 9.7 |
| *Cranoglanis boudierius* | 0.88 | 47.96 | 24,347 | 97.6 | 96.6 | 1 | 0.5 | 1.9 |
| *Danio rerio* | 1.68 | 58.62 | 28,784 | 94.5 | 90.8 | 3.7 | 2.5 | 3 |
| *Hemibagrus wyckioides* | 0.79 | 35.13 | 25,923 | 96.2 | 94.9 | 1.3 | 0.7 | 3.1 |
| *Ictalurus furcatus* | 0.82 | 42.1 | 25,168 | 96.4 | 95.1 | 1.3 | 0.6 | 3 |
| *Ictalurus punctatus* | 0.84 | 43.89 | 26,334 | 97.5 | 96.5 | 1 | 0.4 | 2.1 |
| *Ompok bimaculatus* | 0.72 | 32.16 | 24,575 | 87.1 | 84.6 | 2.5 | 5 | 7.9 |
| *Oreochromis niloticus* | 1.00 | 40.30 | 33,095 | 98.1 | 97.3 | 0.8 | 0.8 | 1.1 |
| *Pangasianodon gigas* | 0.83 | 40.34 | 24,798 | 96.2 | 95.1 | 1.1 | 1.5 | 2.3 |
| *Pangasianodon hypophthalmus* | 0.76 | 36.8 | 25,009 | 97.5 | 96.5 | 1 | 0.8 | 1.7 |
| *Pangasius djambal* | 0.84 | 41.64 | 24,372 | 94.3 | 93.3 | 1 | 2.5 | 3.2 |
| *Silurus aristotelis* | 0.80 | 41.87 | 23,628 | 96.8 | 95.7 | 1.1 | 0.8 | 2.4 |
| *Silurus asotus* | 0.75 | 34.55 | 24,039 | 95.4 | 93.9 | 1.5 | 1.8 | 2.8 |
| *Silurus glanis* | 0.79 | 33.91 | 23,905 | 86 | 84.1 | 1.9 | 5.7 | 8.3 |
| *Silurus meridionalis* | 0.74 | 38.5 | 25,402 | 94.3 | 92.8 | 1.5 | 1.6 | 4.1 |
| *Tachysurus fulvidraco* | 0.71 | 35.14 | 24,013 | 96.7 | 95.5 | 1.2 | 0.7 | 2.6 |
| *Tachysurus vachellii* | 1.04 | 33.26 | 23,735 | 96.7 | 95.7 | 1 | 0.9 | 2.4 |
| *Trichomycterus rosablanca* | 0.69 | 55.03 | 21,171 | 96.3 | 95 | 1.3 | 0.9 | 2.8 |

**Supplementary Table 4:**
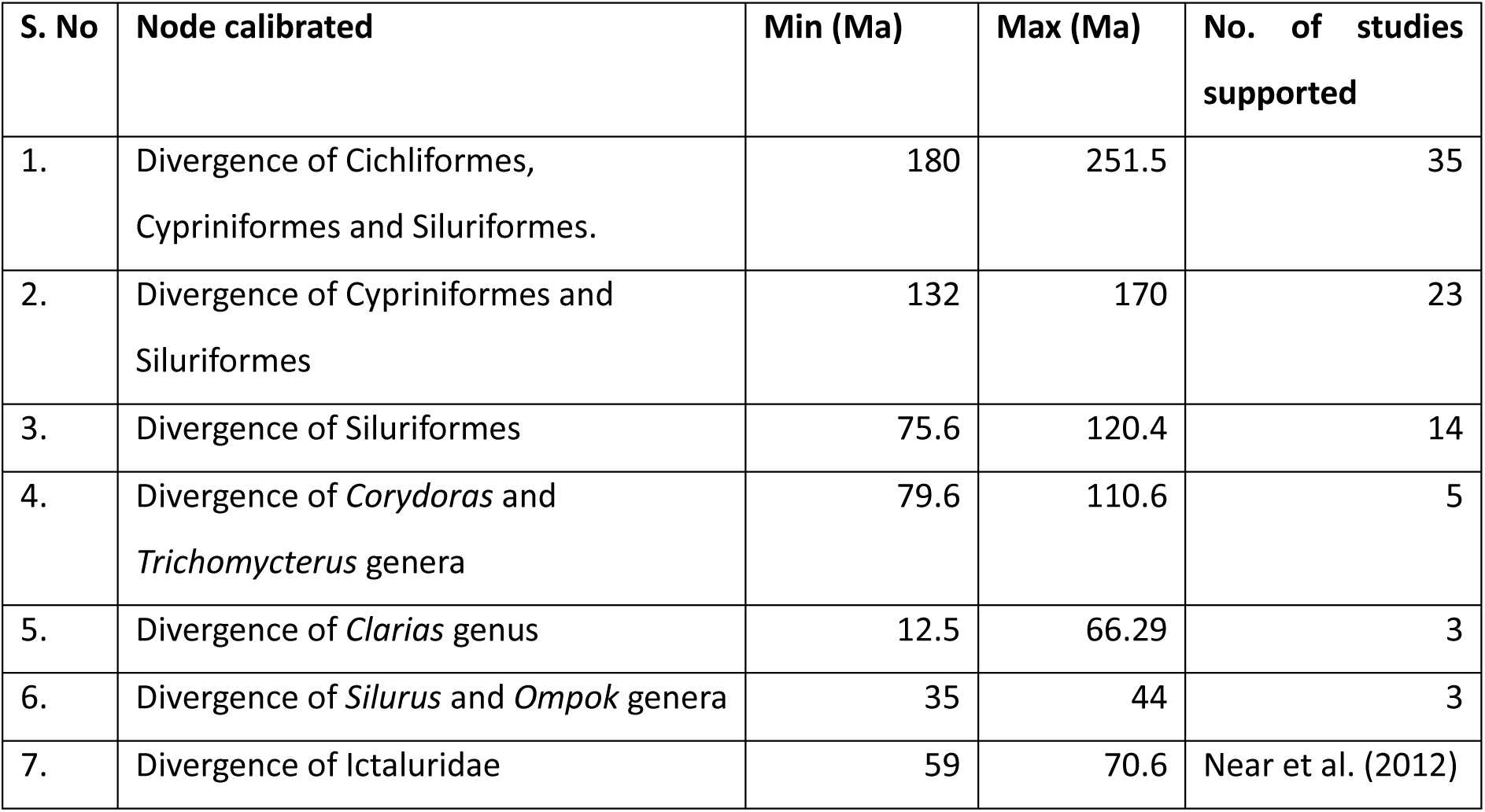
Calibration points for divergence time estimation.

| S. No | Node calibrated | Min (Ma) | Max (Ma) | No. of studies supported |
| --- | --- | --- | --- | --- |
| 1. | Divergence of Cichliformes, Cypriniformes and Siluriformes. | 180 | 251.5 | 35 |
| 2. | Divergence of Cypriniformes and Siluriformes | 132 | 170 | 23 |
| 3. | Divergence of Siluriformes | 75.6 | 120.4 | 14 |
| 4. | Divergence of *Corydoras* and *Trichomycterus* genera | 79.6 | 110.6 | 5 |
| 5. | Divergence of *Clarias* genus | 12.5 | 66.29 | 3 |
| 6. | Divergence of *Silurus* and *Ompok* genera | 35 | 44 | 3 |
| 7. | Divergence of Ictaluridae | 59 | 70.6 | Near et al. (2012) |

